# Efficiency of RNAi based gene silencing in fungi - a review and meta-analysis

**DOI:** 10.64898/2026.03.04.709060

**Authors:** Patrick Barth, Jakob Drumm, Alexandra Schmidt, Florian Hartig, Aline Koch

## Abstract

RNA interference (RNAi) shows great potential to protect crops against fungal diseases, yet reported protection efficiencies vary greatly, and our understanding of the factors responsible for this variance remains limited. In this meta-analysis, we evaluated 89 studies that compare the efficiency of host-induced gene silencing (HIGS) and spray-induced gene silencing (SIGS) in controlling fungal diseases, focusing on biotrophic, hemibiotrophic, and necrotrophic fungi, the use of formulations, and the dsRNA design as explanatory factors for differences between reported efficiency values. Our results indicate that SIGS is slightly more effective, particularly in biotrophs. Surprisingly, SIGS studies using formulations did not outperform those applying naked dsRNA. We also assessed parameters of RNA design. Differences in dsRNA length and the number of constructs, and number of targets showed no consistent significant effect on resistance in either HIGS or SIGS. Interestingly, however, HIGS studies reported significantly higher efficiency when targeting genes closer to the 3’ end and SIGS when targeting genes closer to the 5’ end. We discuss potential reasons for the reported patterns, such as variability in dsRNA uptake mechanisms, intercellular trafficking and Dicer processing, and conclude that more research is needed to understand the biological mechanisms determining RNAi efficiency for fungal control.

## Introduction

RNA interference (RNAi) is a process that occurs throughout eukaryotic life where small RNA molecules can silence specific gene expression. RNAi was first discovered as sequence-specific co-suppression in plants (Napoli et al. 1990) but was subsequently identified in animals as well. Fire and Mello received the Nobel Prize for their pioneering work in this area (Fire et al. 1998). RNAi can interfere with gene expression at the levels of transcription and translation. It starts when cellular double-stranded RNAs (dsRNAs), which can originate from viral replication or transposable elements, are processed by DICER-LIKE RNases (DCLs) into small RNAs (sRNAs). These small interfering RNAs (siRNAs) are incorporated into the RNA-induced silencing complex (RISC) by loading onto ARGONAUTE (AGO) proteins. Within this complex, the guide strand of the siRNA directs the recognition of complementary messenger RNA (mRNA), leading to target cleavage, degradation, or translational repression. This process, collectively termed post-transcriptional gene silencing (PTGS), is mediated by conserved proteins across plants, fungi, and animals (Filipowicz et al. 2005; Ashfaq et al. 2020; Cogoni and Macino 2000)(Filipowicz et al. 2005; Ashfaq et al. 2020; Cogoni und Macino 2000). After the discovery of RNAi, researchers soon realised its many potential biotechnological applications. Medical applications, for example, use RNAi to inhibit the synthesis of harmful proteins (Fattal und Bochot 2006; Hu-Lieskovan et al. 2005). A particularly interesting and novel application is phytomedicine, where RNAi could be used to protect plants against diseases caused by phytopathogens and pests. The first studies based on this idea used transgenic plants that express double-stranded RNA (dsRNA) (Nowara et al. 2010; Koch et al. 2013). This approach, called host-induced gene silencing (HIGS), was applied in the first transgenic RNAi corn variety, ‘SmartStaxPro’, approved in 2018 (Head et al. 2017)210–240 nucleotide (nt) dsRNA targeting the. Numerous studies have since confirmed the efficacy of HIGS against economically important pathogens and pests.

Although clearly effective, the proliferation of RNAi-based crop protection via HIGS is hindered by three factors: (i) the technical challenges and thus lengthy timelines required for stable host plant transformation, (ii) associated high development and regulatory costs, estimated at up to $140 million per transgenic crop (Rosa et al. 2018), and (iii) ongoing global public resistance to genetically modified crops (Tardin-Coelho et al. 2025). The main alternative that is therefore pursued is to apply dsRNAs exogenously to plants, known as spray-induced gene silencing (SIGS).

The efficacy of the SIGS approach was first demonstrated by (Koch et al. 2016), who showed that foliar application of dsRNAs provides effective protection against the necrotrophic fungal pathogen *Fusarium graminearum*. Since then, multiple investigations have corroborated the efficacy of SIGS against a wide range of plant diseases (Qiao et al. 2021; S et al. 2022; Spada et al. 2023; Rank und Koch 2021). In 2023, the first SIGS-based product (Calantha^TM^) was approved by the US Environmental Protection Agency (EPA). It provides protection against the Colorado potato beetle (CPB) via foliar application to potato leaves (Yan et al. 2024).

Compared to HIGS, SIGS has several practical advantages: it can be deployed more quickly and cheaply, and it can be used on a wider range of crop species, as it is not limited by plant transformability (Uslu et al. 2025). However, it has also been hypothesized that SIGS may confer lower and less durable resistance than HIGS, as the latter results in substantially higher and sustained dsRNA levels due to continuous transgene expression rather than external application, which is prone to environmental degradation and must overcome multiple uptake barriers. This hypothesis, however, has not yet been formally tested. Interestingly, direct comparisons of HIGS and SIGS in controlled experiments have shown that sprayed RNAs can sometimes be more effective than transgene-based expression (Hu et al. 2020; Koch et al. 2019). The superior performance of SIGS in some cases may be explained by the ability of necrotrophic fungi to take up unprocessed dsRNA (Wang et al. 2016; Qiao et al. 2021; Brosnan et al. 2021), which is then processed by their own fungal RNAi machinery (Koch et al. 2016) (Fig. 1A).

**Fig. 1:**
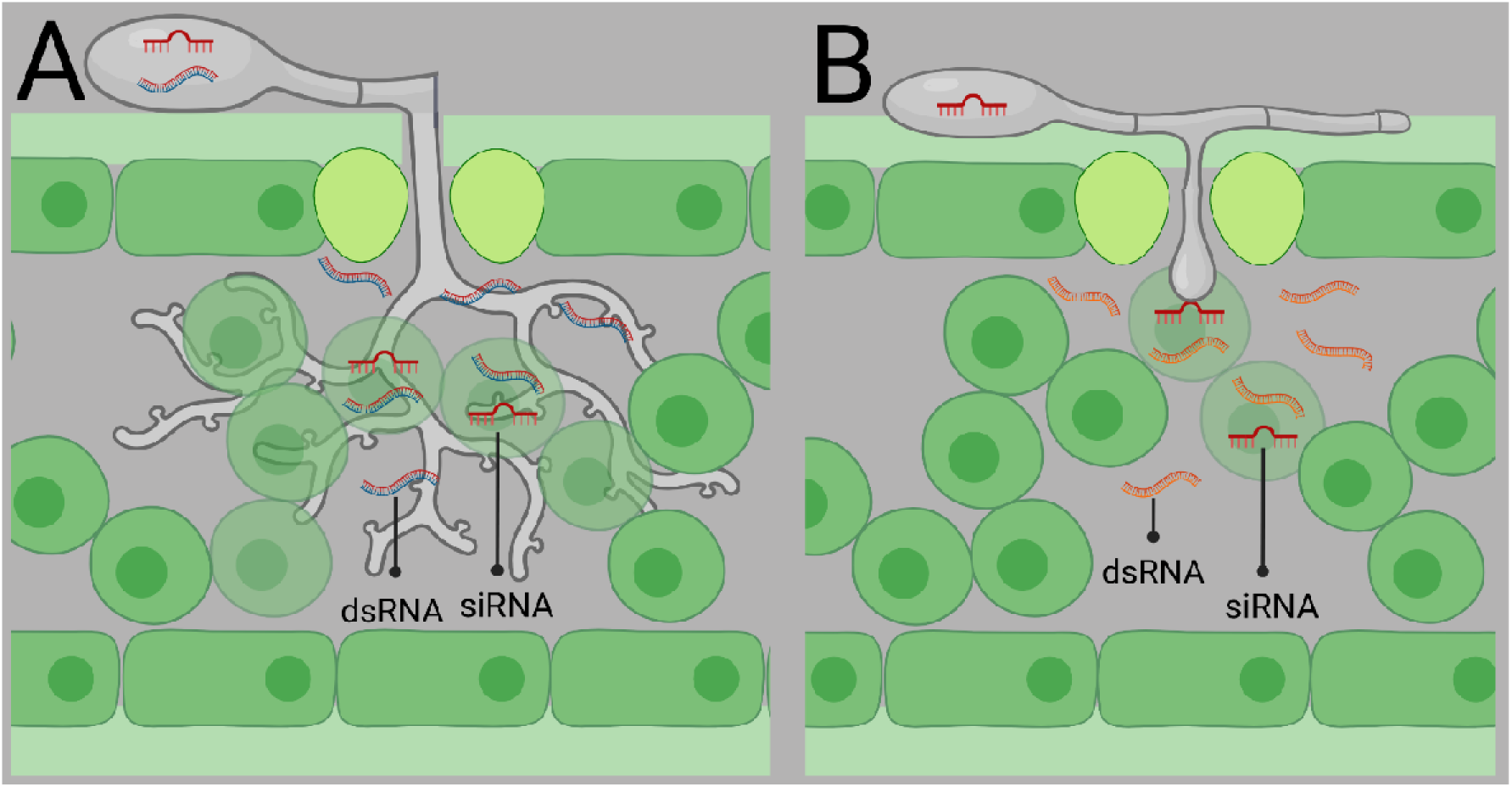
Proposed model linking fungal lifestyle to RNA uptake and processing mechanisms. Based on the findings (from this and other studies) that some necrotrophic fungi can take up unprocessed dsRNA, probably from the apoplast, and process it into siRNAs using their own RNAi machinery (A). This potentially results in more “effective” siRNAs being produced directly within the target organism (fungus) and thus presumably cause a greater effect. In contrast, a biotrophic fungus relies on the uptake of cellular siRNAs from the host plant, i.e., siRNAs that have previously been processed from transgene-derived dsRNAs by the plant’s RNAi machinery (B). This implies that, in SIGS, multiple barriers, including the plant cell wall, must be overcome before siRNAs become accessible for uptake by a biotrophic fungus. Therefore, HIGS may be a more effective delivery strategy for controlling biotrophs, as it ensures a continuous supply of plant-derived siRNAs.

Although both approaches, HIGS and SIGS, rely on the same PTGS mechanism, they differ fundamentally in the origin and processing of dsRNA. In HIGS, dsRNA is generated endogenously and processed by plant DCL enzymes into siRNAs, which can be delivered to the pathogen, possibly via extracellular vesicles (EVs) (Fig. 1A) (Wang et al. 2024b). In contrast, SIGS involves the exogenous application of dsRNA, which must cross plant structural and cellular barriers prior to uptake by the pathogen (Fig. 1B). Based on these mechanistic distinctions and given reports that necrotrophic fungi can internalize and process unprocessed dsRNA, while biotrophic fungi are less exposed to extracellular RNA pools, we hypothesize that necrotrophic fungi could be more efficiently targeted via SIGS, whereas control of biotrophic fungi would benefit more from the constitutively produced plant-derived cellular siRNAs in HIGS. To evaluate whether the current literature supports this working model, we performed a meta-analysis on all studies that reported RNAi efficiency against fungal phytopathogens. We first performed a systematic search using PubMed, complemented by manual searches in other scientific databases, which resulted in 114 published studies, of which 89 met our inclusion criteria for the final analysis. We extracted and standardized (e.g., by normalizing and converting to comparable metrics) the reported RNAi efficiency. We also extracted all factors that could potentially affect RNAi efficiency, such as delivery method, fungal lifestyle, or dsRNA length. We then performed a meta-analysis based on these data, addressing the following research questions: i) How does RNAi efficiency compare between the two major delivery methods, HIGS and SIGS, controlling for all other experimental factors? ii) How does RNAi efficacy differ between necrotrophic and biotrophic fungi, given their distinct lifestyles? iii) How does dsRNA length and structure influence RNAi efficiency? Our results show several patterns that are both interesting for obtaining a better understanding of the biological mechanisms of RNAi and also provide guidance for developing RNAi applications for disease control.

## Material and Methods

### Data collection for the meta-analysis

For the meta-analysis, we queried the PubMed database [https://pubmed.ncbi.nlm.nih.gov/] on August 5, 2024, for all publications that applied HIGS or SIGS in the context of plant resistance against fungal pathogens (for the exact search terms, see Supporting Method S1). This resulted in 72 studies. We identified 42 additional publications through manual literature search. Altogether, this resulted in 114 unique publications (Table S1).

All 114 publications were evaluated according to predefined inclusion criteria (see Supporting Method S2, section “Inclusion criteria for publications”). Briefly, publications were excluded if they did not investigate plant resistance against fungal pathogens, did not examine the effects of HIGS or SIGS, or did not report results that could be expressed as a numerical value of resistance transfer. As a result, 25 publications were excluded, resulting in 89 studies that met the predefined inclusion criteria (see Supporting Method S2, section “Study selection”) for the meta-analysis.

From these 89 studies, we extracted 468 distinct measurements of pathogen resistance after RNAi application (see Supporting Method S3). Resistance was defined as the relative reduction of disease symptoms in treated plants compared to untreated controls. Where available, we extracted resistance measurements and standard errors as numeric values directly from the text or tables. If resistance values and standard errors were not explicitly reported, they were calculated based on the measurements of the control and treatment groups (see Supporting Method S4, section “Calculation of resistance and error propagation”). In cases where results were presented only in figures without providing numerical values, we used the R package *metaDigitise* (Pick et al. 2019) to extract resistance values and standard errors from the figures.

In addition to the resistance values, we extracted a range of experimental details from each study (for details see Supporting Method S4, section “Data extraction”). This included the taxonomic classification of plant and fungal species, the fungal lifestyle, target gene information (e.g., gene ID and CDS length), dsRNA characteristics (e.g. dsRNA length and position within the target gene), the number of RNAi constructs and target genes, the applied RNAi method (HIGS or SIGS). For SIGS experiments, we additionally evaluated whether a formulation was used and what dsRNA concentration was used. Resistance measurement methods were summarized into five categories: general reduction, infected leaf area, biomass, haustoria index, and sporulation (for details see Supporting Method S2, section “Inclusion criteria for publications”). Experiments that did not explicitly specify if a formulation was used were assigned to the group “unknown”. A full list of extracted variables is available in the supplements (Supporting Method S4)

### Data preparation

For single experiments with multiple values (e.g., target gene length or dsRNA length in experiments targeting multiple genes or employing multiple dsRNAs), we summarized these values by their mean to avoid overrepresenting these results. This also applied to the dsRNA start and end positions on the target gene(s).

Some publications reported multiple experiments that were performed under similar conditions, differing only in minor aspects (e.g. dsRNA concentration or measurement method). To avoid overrepresenting these experiments, they were either excluded or merged into a single representative entry. Experiments were considered similar when they shared the exact same RNAi method, fungal and plant species and strain, target gene characteristics, number of constructs and target genes, dsRNA position on the target gene, assay type, formulation, and measurement method, while originating from the same publication. For 42 experiments that differed only in dsRNA or formulation concentration, the entry with the highest resistance measurement was retained, while the others were excluded, resulting in 12 representative entries. The reason for this was to remain consistent with the DPI selection, where only the highest resistance value was selected for data extraction (see Supporting Method S3). For 177 experiments, that differed only in the measurement method, the mean resistance value and standard error were calculated, resulting in 82 representative entries. After this merging procedure, we obtained a final dataset of 343 resistance values.

### Data imputation and scaling

In some publications, certain variables that were selected for extraction were missing. Of the 343 entries (comprising 2,401 data points), 240 contained at least one missing value, resulting in a total of 436 missing data points. Since linear regression models require complete data for all included predictors, we used the R package missRanger (Mayer 2024) to impute missing values for standard error, dsRNA length, formulation, target gene CDS length, relative coverage of the target gene by the dsRNA, and the number of constructs and target genes. The imputation was based on all named predictors, in addition to the resistance value, RNAi method, and fungal class and lifestyle. After imputation, numerical predictors were scaled and centered to ensure comparability of effects across all predictors.

### Overview of the experimental data

In the resulting dataset, the number of experiments performed on biotrophic and necrotrophic fungi was very similar (133 and 152, respectively), whereas hemibiotrophic fungi were underrepresented, with only 58 experiments included. When further dividing the experiments by RNAi method, HIGS and SIGS were applied in comparable numbers to biotrophic (75 vs. 58) and necrotrophic fungi (72 vs. 80). In contrast, RNAi experiments performed on hemibiotrophic fungi mainly employed HIGS (48 vs. 10). Notably, only one SIGS experiment reported a resistance value below 13.75%, representing the lowest resistance value observed (−28.17%). In contrast, 10 HIGS experiments reported resistances below that value, with two reducing the plant resistance compared to the control, resulting in negative resistance values, and three showing no change in resistance (Fig. 2).

**Fig. 2:**
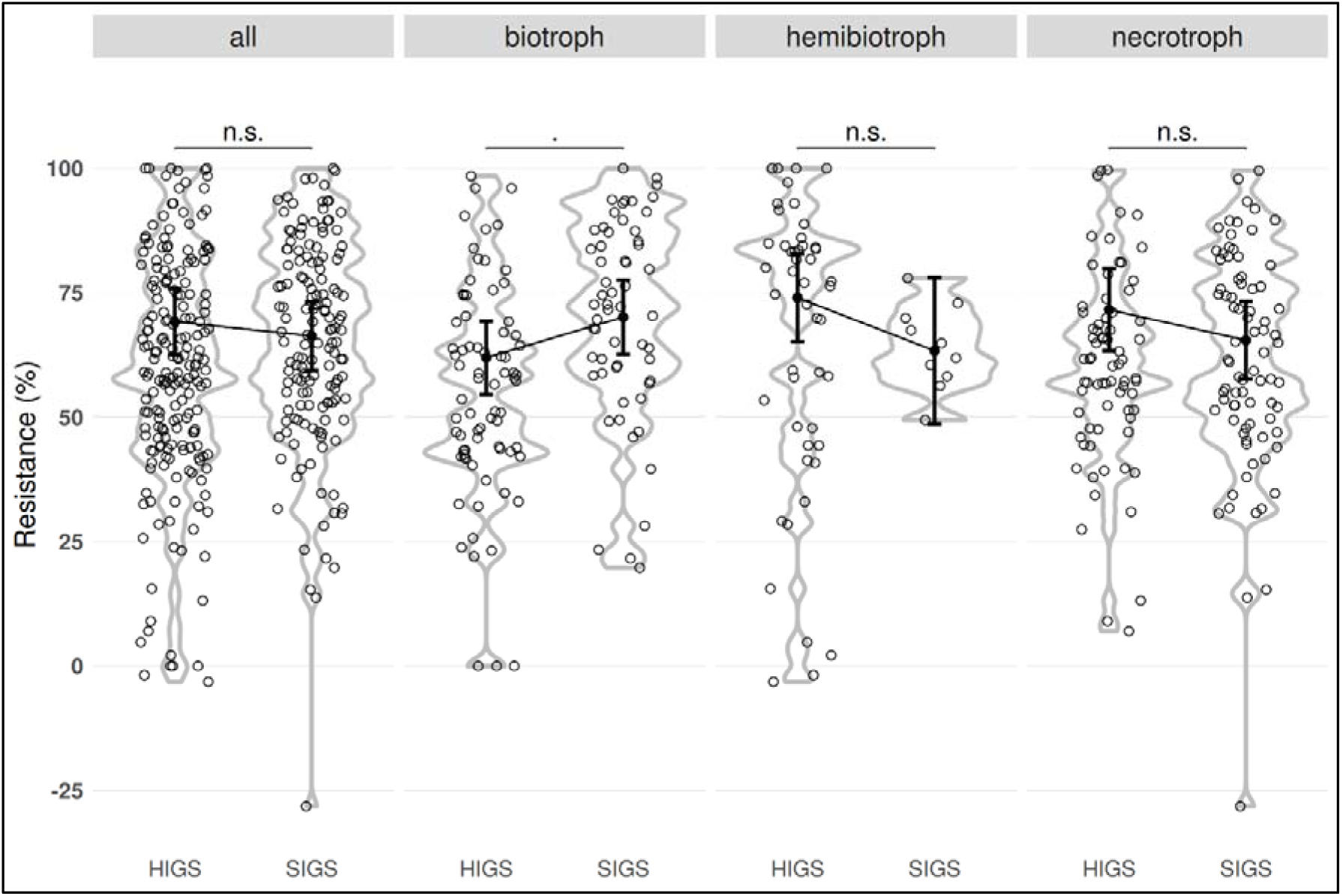
Difference in resistance between RNAi methods overall and grouped by fungal lifestyle, unadjusted for other explanatory variables. Each point represents a distinct experiment. Displayed means and error bars represent estimated marginal means and 95% confidence intervals. Significance levels are based on pairwise comparisons from the marginal means analysis: * p<0.05; ** p<0.01; *** p<0.001. Note that all displayed effects and p-values are marginal, i.e., not adjusted for other covariates.

### Meta-regression

To identify which factors predict resistance, we performed a hierarchical meta-regression with resistance (res) as the response variable, an observation-level random effect to account for biological variability between experiments, experimental standard errors ε_obs_ extracted from the studies, and fungal lifestyle and class, dsRNA length, number of constructs and target genes, and the relative dsRNA position on the target gene (for details, see Supporting Method S4, section “Calculate relative position of dsRNA on target gene”), transfection, and formulation as predictors, each interacting with the RNAi method. The linear regression model was fitted using the *lmer* function from the R package lme4 (Bates et al. 2015) together with the R package *lmerTest* (Kuznetsova et al. 2017). The full structure of the fitted model was:

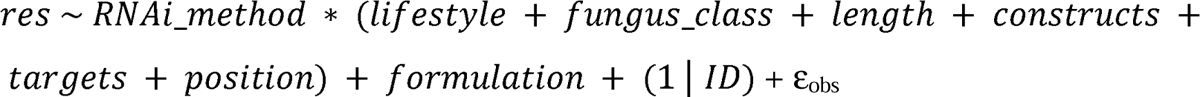

To evaluate the separated contribution of individual predictors, we calculated Type II ANOVA R^2^ contributions using the anova.lmerModLmerTest method from the R package lmerTest. The total R^2^ for the fixed effects was calculated using the *r.squaredGLMM* function from the R package MuMIn (Bartoń 2025). The R^2^ for every separate fixed effect was calculated based on the sum of squares (SUM Sq) from the ANOVA results and the total R^2^ of the fixed effects using the following formula:

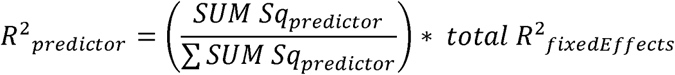

### Marginal effects and visualization of results

The meta-regression estimates the effect of each predictor while all other predictor variables are corrected for, thus implicitly being held at a value of zero. As we included various interactions, this effect is not necessarily identical to the visual effect as assessed from a scatter plot of resistance against the predictor variable, which corresponds to the marginal (or average) effect of the predictor while the other variables vary across their observed ranges. To display the estimated effects of the meta-regression together with scatter plots of the data, we therefore calculated marginal mean effects, their 95% confidence intervals, and corresponding p-values using the *emmeans* function from the R package emmeans (Lenth 2025).

## Results and Discussion

### SIGS provides higher overall resistance while HIGS is more effective against necrotrophic fungi

Although HIGS provides a continuous intracellular source of dsRNA and bypasses the multiple uptake barriers that exogenous RNA must overcome, our meta-analysis did not find that it is overall more effective than SIGS. When looking at the marginal comparison of all HIGS and SIGS values without correcting for other explanatory factors, we did not find any significant effect. When comparing HIGS and SIGS separately for the three fungal lifestyles, but not correcting for other factors, we find that SIGS performed significantly better than HIGS against biotrophs, while no significant differences can be found for hemibiotrophs and necrotrophs (Fig. 2). When adjusting for all covariates in the meta-regression, our results indicate that experiments using SIGS resulted in significantly higher resistance (Table 2).

Looking at the resistance differences between fungal lifestyles for either HIGS or SIGS, unadjusted for other covariates, we find that HIGS provides significantly higher resistance in hemibiotrophs and necrotrophs compared to biotrophs (Fig. 3, as already visible in Fig. 2). These effects also remain after adjusting for all other covariates in the meta-regression (Table 2). Unadjusted for other covariates, studies using SIGS did not show significant differences between the different fungal lifestyles (Fig. 3). However, when adjusting for other covariates in the meta-regression, we find that hemibiotrophs and necrotrophs are significantly less affected by SIGS compared to biotrophs (Table 2).

**Fig. 3:**
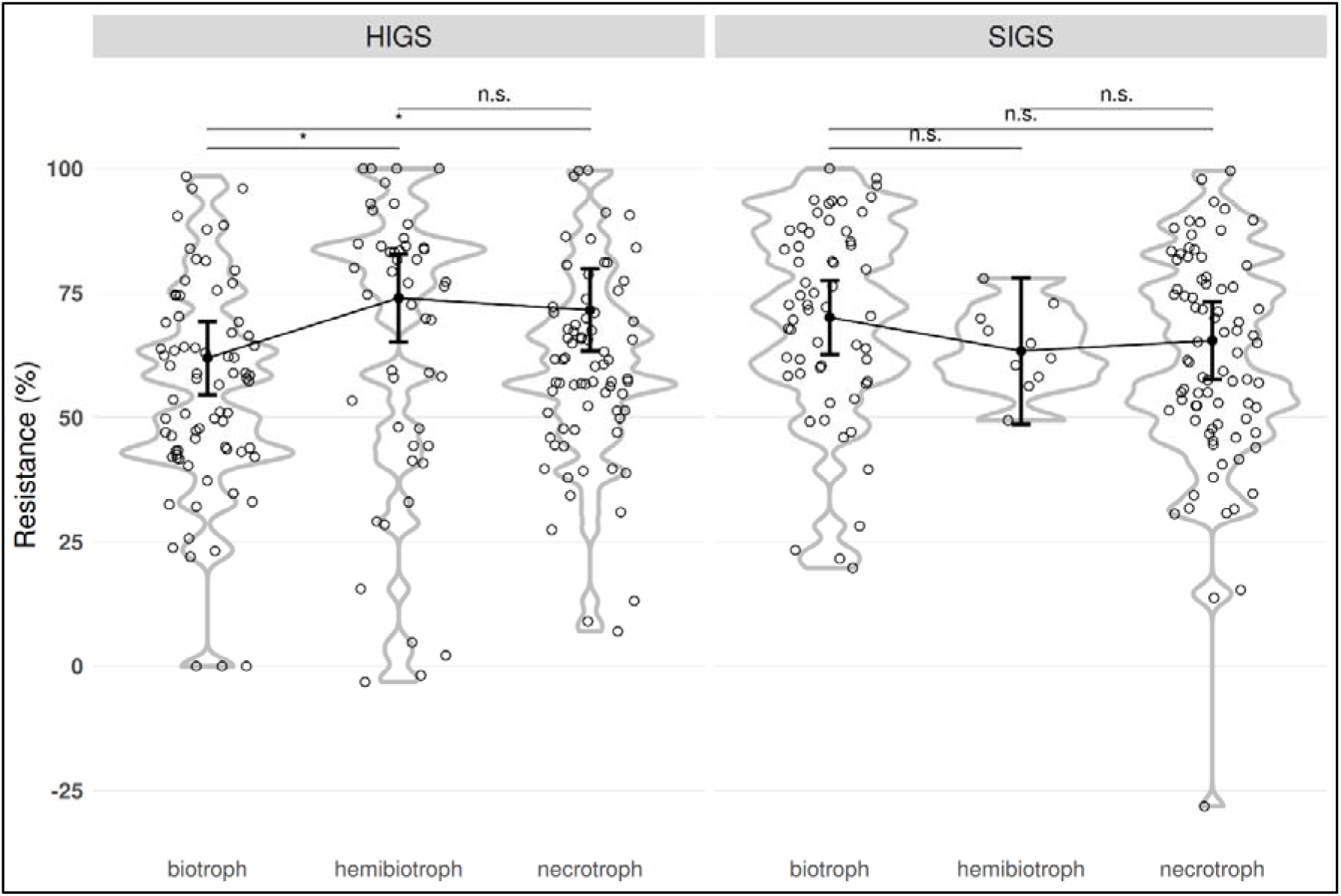
Difference in resistance between fungal lifestyle for HIGS and SIGS, unadjusted for other explanatory variables. Each point represents a distinct experiment. Displayed means and error bars represent estimated marginal means and 95% confidence intervals. Significance levels are based on pairwise comparisons from the marginal means analysis: * P<0.05; ** P<0.01; *** P<0.001. Note that all displayed effects and p-values are marginal, i.e., not adjusted for other covariates.

Overall, these results suggest that differences in RNA uptake or processing between fungal lifestyles may contribute to the varying effectiveness of both RNAi methods and that experiments based on SIGS tend to achieve higher resistance, particularly for biotrophs. We carefully checked the data to see if these results could be affected by experimental errors. The lowest measured resistance in SIGS was –28.17%, likely an outlier caused by the LDH (layered double hydroxide) formulation used in the experiment (see Fig. 6C in (Mukherjee et al. 2024)). LDH is a formulation that protects the dsRNA from environmental conditions, prevents wash-off, and enables its slow release from the matrix (Mitter et al. 2017). The second-lowest SIGS measurement was 13.75% (Wu et al. 2024). In contrast, 10 HIGS experiments reported resistance values below 13.75%. A comparison of raw mean resistance values also suggests that SIGS outperforms HIGS, with mean resistance levels of 65.1% for SIGS and 59.4% for HIGS. Several publications featured in the meta-analysis performed both HIGS and SIGS experiments under comparable conditions: Hu et al. investigated the resistance of soybean plants (*Glycine max*) against soybean rust (*Phakopsora pachyrhizi*) after HIGS and SIGS treatments, observing a greater reduction in pustule density and fungal biomass with SIGS (Hu et al. 2020). Koch et al. examined the inhibition of *Fusarium graminearum* growth on barley plants (*Hordeum vulgare*) after HIGS and SIGS using different constructs targeting *FgCYP51* genes. SIGS showed higher resistance with all constructs used (Koch et al. 2019). Altogether, we believe that the empirical finding that experiments using SIGS report higher effectiveness than experiments using HIGS is robust.

These results are somewhat in contradiction to our theoretical model, which had predicted higher efficacy of SIGS primarily in necrotrophic fungi, as they can directly absorb and process externally applied dsRNA (Koch et al. 2016; Qiao et al. 2021; Brosnan et al. 2021). Based on our analysis, we speculate that biotrophic and hemibiotrophic fungi may also be capable of absorbing high amounts of RNA, especially during the early infection stages. Several pieces of evidence support this possibility: First, successful HIGS experiments against obligate biotrophs, such as powdery mildews and rusts, demonstrate that siRNAs and/or dsRNAs can be transferred from host to pathogen. Second, haustoria, the feeding structures formed by many biotrophs, represent intimate interfaces that may facilitate the uptake of plant-derived RNA. Third, it has been shown that plants export small RNAs via EVs into interacting fungi. This process has been demonstrated for both necrotrophic and biotrophic lifestyles (Cai et al. 2018). These mechanisms offer a plausible explanation as to why SIGS can sometimes match or even surpass HIGS, even in biotrophic systems. Another possible explanation for our results could be differences in the effective dsRNA exposure of pathogens. The rapid uptake of dsRNAs during the early infection stages may enhance the efficiency of SIGS, as fungal spores and germ tubes are directly exposed to RNA molecules prior to host tissue penetration. Furthermore, following spray application, the concentration of dsRNAs in the upper epidermal tissues is likely to be higher than in HIGS, especially around stomata and trichomes (Koch et al. 2016), which represent natural entry sites for many biotrophic fungi. This early contact increases the likelihood of silencing essential fungal genes at critical developmental stages, e.g., during sporulation or germ tube formation, which may enhance RNAi efficiency before haustoria are established, thereby limiting pathogen colonization and subsequent disease progression. This effect could be further enhanced if the fungus acquires both unprocessed dsRNA and plant-processed siRNAs. While biotrophs secrete effectors into the apoplast in order to manipulate the host organism (Ellis et al. 2009), this could also work bidirectionally, leading to the uptake of dsRNAs present in the apoplast. However, HIGS has the advantage of continuous dsRNA production, resulting in sustained resistance effects, whereas SIGS requires repeated applications to maintain its efficacy.

The higher effect of HIGS against necrotrophs could be explained by their potentially higher exposure to environmental as well as plant-derived RNAs due to the active degradation of host tissue, compared to biotrophs, which maintain living host cells. To test this hypothesis, targeted experiments would be required. For example, plant *dcl* mutants, which mainly produce unprocessed dsRNA rather than siRNAs, could be infected with biotrophic and necrotrophic fungi to examine whether differences in uptake or processing could help to explain the observed levels of resistance. Another possible explanation is the recently described export of endogenous plant RNAs to interacting necrotrophic fungi via EVs and other secretion pathways (Wang et al. 2024a). If necrotrophs encounter and internalize more of these exported RNAs due to their feeding strategy, they could receive higher effective doses of silencing RNAs than biotrophs. Further investigation of these alternative mechanisms is needed to clarify whether the higher effectiveness of HIGS against necrotrophs reflects differences in fungal uptake, the form of RNA delivered (dsRNA vs. siRNA), or plant export dynamics.

### Formulation shows no apparent improvement of effectiveness in SIGS

Our third hypothesis was that the use of a formulation in SIGS experiments increases effectiveness compared to naked dsRNA. Additionally, we hypothesized that naked dsRNA has a higher efficiency against necrotrophic fungi than biotrophic fungi. Neither the raw correlations, unadjusted for other covariates (Fig. 4A), nor the meta-regression adjusted for all other covariates (Table 2) showed a significant difference between naked RNA and the use of a formulation. This non-significant result also remained when comparing the effect of naked RNA across the different fungal lifestyles (Fig. 4B).

**Fig. 4:**
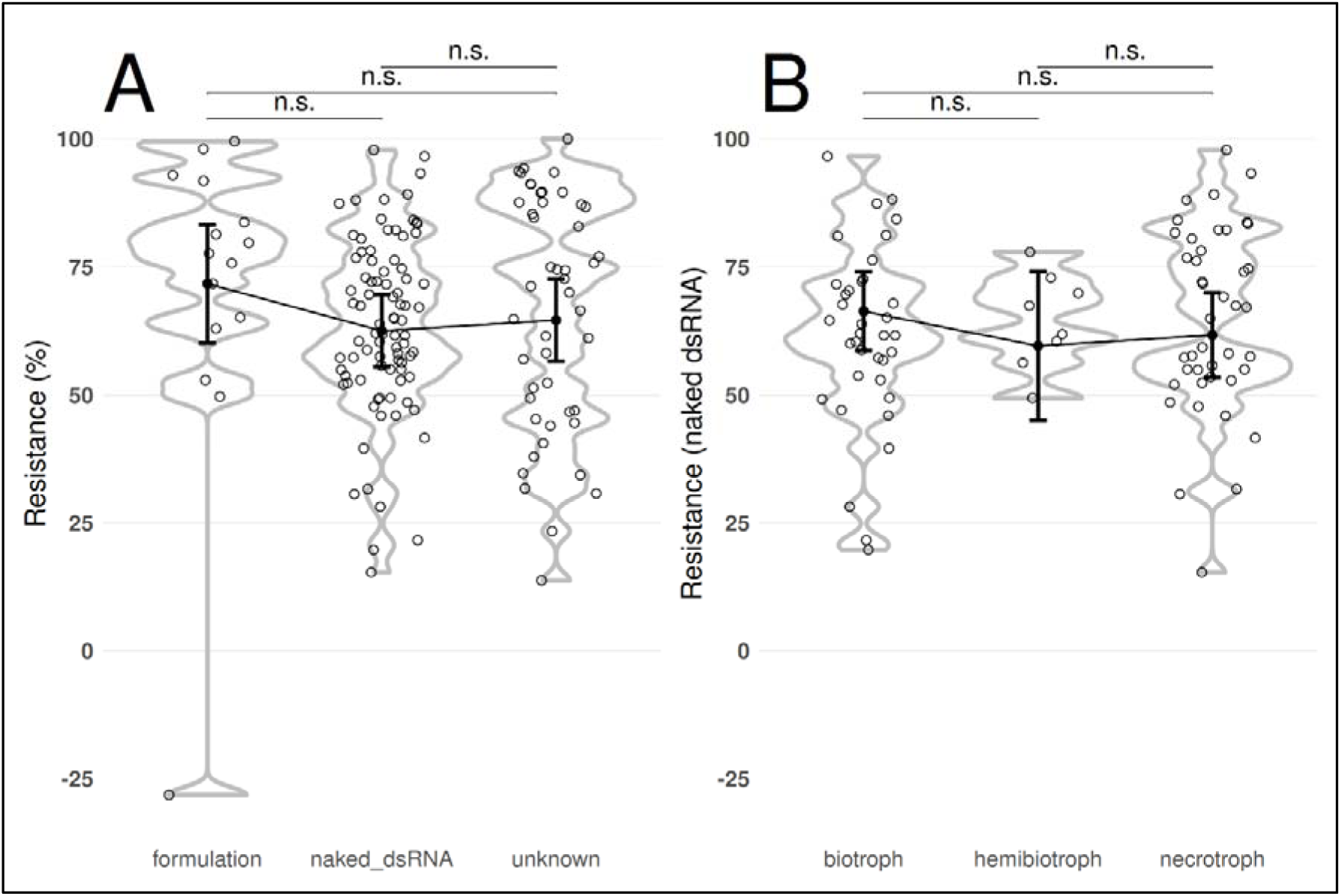
Effect of formulation on resistance in SIGS experiments, unadjusted for other explanatory variables. Due to the high variability among formulations, all experiments involving a formulation were grouped together. Furthermore, several publications did not specify whether a formulation or naked RNA was used and were therefore categorized as “unknown”. (A) Additionally, experiments employing SIGS with naked dsRNA were further divided by fungal lifestyle (B). Each point represents a distinct experiment. Displayed means and error bars represent estimated marginal means and 95% confidence intervals. Significance levels are based on pairwise comparisons from the marginal means analysis: * P<0.05; ** P<0.01; *** P<0.001. Note that all displayed effects and p-values are marginal, i.e., not adjusted for other covariates.

In interpreting these results, we note that a lack of formulation effects on experimental resistance values does not necessarily mean that formulations offer no other advantages. Some formulations have been shown to enhance DNA uptake into plant cells (Demirer et al. 2019) or increase the longevity of dsRNA molecules by protecting them from environmental conditions (Mitter et al. 2017). This may not lead to increased resistance shortly after application but rather to a more prolonged effect over time. Such sustained effects were demonstrated in studies by Niño-Sanchez et al. and Qiao et al., both of which reported only minor differences in the applied resistance when comparing the formulation with naked dsRNA at 1 dpi. However, when resistance was observed over longer periods, it became evident that the effect of naked RNA diminished considerably faster compared to samples in which a formulation was used (Niño-Sánchez et al. 2022; Qiao et al. 2023). Due to our decision to include only one measurement per experiment, this temporal effect was not captured in our meta-analysis.

Moreover, our own results and those of others indicate that whether naked RNA can be taken up efficiently depends on the plant species. Thus, for some plants, a formulation may not be necessary, or indeed could be disadvantageous, since the RNA would then need to be released from the carrier matrix, which may not occur with full efficiency. This is particularly relevant because most of the included studies were conducted under controlled conditions rather than in the field, where the stability advantage of formulations would be more relevant.

It should also be noted that formulations are often used in experiments where the use of naked RNA alone failed to produce sufficient uptake. However, those initial failed results are usually not reported in the final publications. In other words, formulations are often used to elevate ‘non-responsive’ systems to the level of responsive ones, and one would not necessarily expect a strong overall difference across the final study results.

### dsRNA length has no significant impact on resistance

Finally, we addressed hypotheses on how dsRNA design may influence the efficacy of HIGS and SIGS approaches. First, we conjectured that the length of dsRNAs does not influence resistance in HIGS experiments, while in SIGS experiments, different lengths may have varying effects on observed resistance, with longer dsRNAs being less effective due to reduced uptake efficiency (Höfle et al. 2020). We also hypothesized that a higher number of constructs and targets would lead to increased efficiency, while the position of the target site within the mRNA would have no effect, provided that the targeted region lies within the coding sequence (CDS). This is because targeting untranslated regions (UTRs) is generally considered less effective (Cedden et al. 2025).

Neither unadjusted effects (Fig. 5A & B) nor the adjusted effects from the meta-regression (Table 2) indicate that dsRNA length has a significant effect on resistance in either RNAi method. However, although not statistically significant, SIGS shows a tendency toward reduced effectiveness with increasing dsRNA length (Fig. 5B). The finding that dsRNA length has no significant effect in HIGS was expected, as the dsRNAs are produced within plant cells and therefore do not need to cross cellular barriers. In contrast, it is surprising that no significant effect was observed in SIGS, especially considering that all SIGS experiments using dsRNAs longer than 1,500 nucleotides resulted in relatively low resistance values ranging from 47.7% to 59.3%. Höfle et al. performed a series of SIGS experiments targeting *FgCYP* genes using dsRNAs of varying lengths, ranging from 400 to 1,575 nucleotides. Their results showed that SIGS becomes less effective beyond a certain dsRNA length. Specifically, dsRNAs of 500 nt were most effective, while resistance decreased at lengths of 800 nt and approximately 1,550 nt (Höfle et al. 2020).

**Fig. 5:**
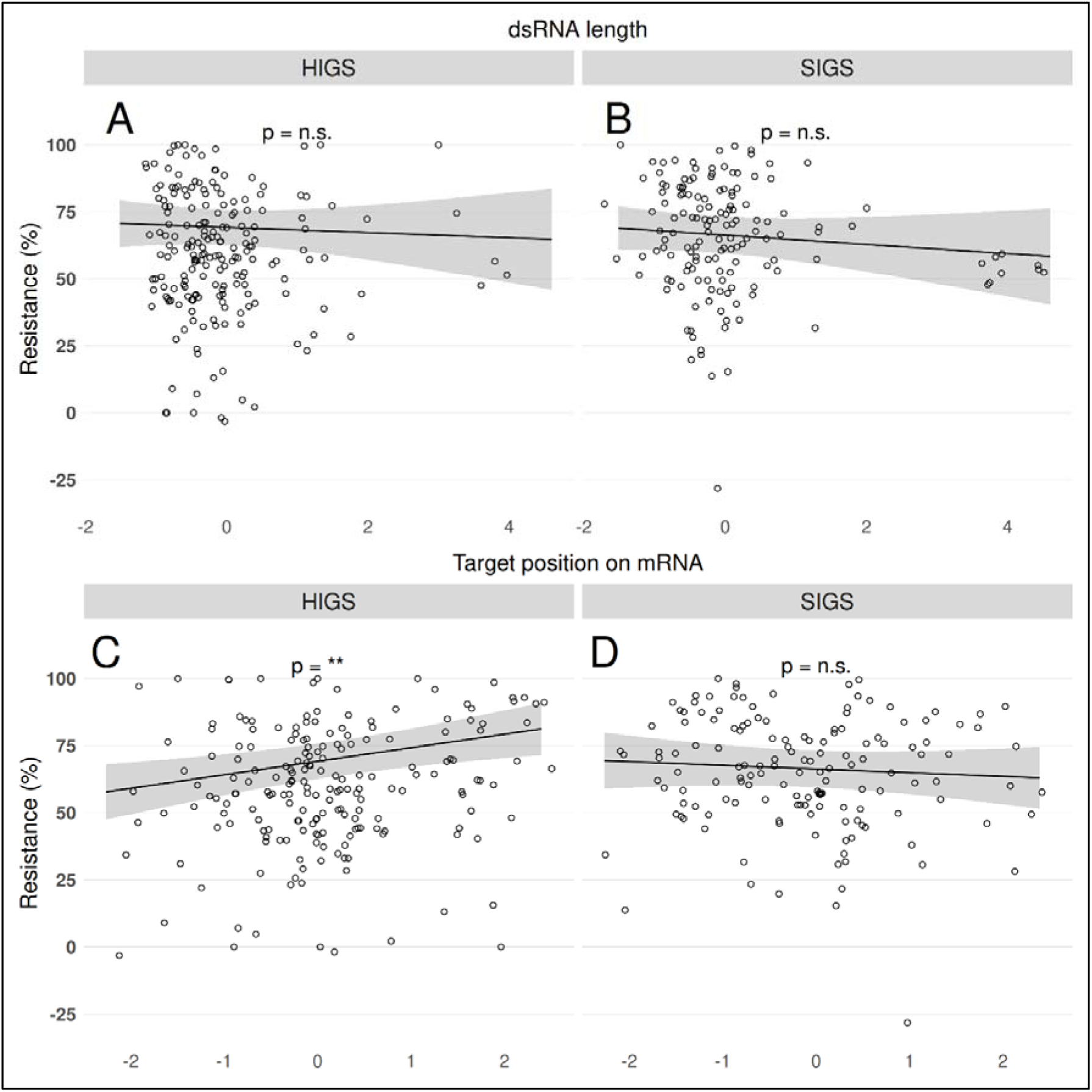
Effect plots of dsRNA design parameters, unadjusted for other explanatory variables. Effect plots showing the impact of dsRNA length in HIGS (A) and SIGS (B), as well as the relative position of the mRNA target site in HIGS (C) and SIGS (D). Each point represents a distinct experiment. The solid line shows the predicted mean and the shaded area the 95% confidence band, both estimated using marginal means. Significance levels are based on the marginal means analysis: * P<0.05; ** P<0.01; *** P<0.001. Note that all displayed effects and p-values are marginal, i.e., not adjusted for other covariates.

Similar tendencies have also been reported by Marais et al. (Marais et al. 2024) in *Cercospora zeina*, where shorter dsRNAs (<800 nt) were more effective than constructs of ∼1.4 kb, and by a recent study on *Botrytis cinerea* showing that a 270 nt dsRNA fragment silenced its target gene more efficiently than a longer chimeric construct (∼470 nt) (Wu et al. 2025). Together, these results support the notion that shorter dsRNAs are more efficiently absorbed and processed during SIGS, while longer dsRNAs may suffer from reduced uptake or delivery efficiency.

The lowered resistance for dsRNAs over 1,500 nt in SIGS is most likely due to more difficult uptake of dsRNAs in plant cells. As externally applied dsRNAs are mostly taken up through stomata (Koch et al. 2016), larger sizes may hinder uptake efficiency.

### HIGS appears to be more effective when targeting regions near the 3’ end

Not adjusting for other covariates, HIGS appears more effective when targeting regions near the 3’ end of the target gene (Fig. 5C), while no comparable effect is observed for SIGS (Fig. 5D). After adjusting for all covariates in the meta-regression, these HIGS effects remain unchanged, whereas an additional effect is observed in SIGS, which appears more effective when targeting regions near the 5’ end of the target gene (Table 2). We speculate that these effects arises due to the accessibility of the mRNA for siRNA binding: regions near the 3’ end might be less structured or contain fewer stable secondary structures related to lower GC content (Wan et al. 2014) than the 5’ end, which can facilitate more efficient RNAi. However, it is important to note that this is not a universal rule, as mRNA structure varies across genes and experimental systems, and some studies may have already selected target sites based on predicted accessibility using RNA design tools. The difference between HIGS and SIGS in relation to target site position may also reflect differences in dsRNA uptake and processing: in HIGS, siRNAs are generated within the plant cells and have direct access to the mRNA, whereas in SIGS, uptake efficiency and processing in the leaf tissue could influence the impact of target site position on RNAi efficiency.

### Number of constructs and targets has no impact on effectiveness in HIGS and SIGS

No significant effect on resistance was observed for the number of constructs or target genes (unadjusted p-values of 0.3538 and 0.7613 for the number of constructs, and 0.1985 and 0.8302 for the number of targets in HIGS and SIGS, respectively). Although the number of constructs and target genes was not the primary focus of our study, we were still surprised to observe no effect at all, as the idea that a higher number of RNAi targets will silence more genes is supported by several studies. For example, Li et al. demonstrated that targeting multiple chitin synthases of rice false smut (*Ustilaginoidea virens*) induced higher resistance than targeting them individually (Li et al. 2021). Another example is provided by Koch et al., who found that multiple targets within the cytochrome P450 genes of *Fusarium graminearum* performed better in both HIGS and SIGS (see Table 1 in (Koch et al. 2019)).

**Table 1:** Number of RNAi experiments included in the meta-analysis, categorized by fungal lifestyle and RNAi method, including mean resistance values and standard errors (SE).

| Lifestyle | Biotroph | Hemibiotroph | Necrotroph | Total |
| --- | --- | --- | --- | --- |
| Sample size all | 133 | 58 | 152 | 343 |
| Sample size HIGS | 75 | 48 | 72 | 195 |
| Sample size SIGS | 58 | 10 | 80 | 148 |
| Mean ± SE | 61.77 ± 1.9 | 64.88 ± 3.48 | 60.78 ± 1.64 | 61.86 ± 1.19 |
| Mean ± SE HIGS | 55.2 ± 2.45 | 65.08 ± 4.18 | 59.98 ± 2.26 | 59.4 ± 1.64 |
| Mean ± SE SIGS | 70.26 ± 2.61 | 63.93 ± 2.67 | 61.49 ± 2.42 | 65.1 ± 1.64 |

**Table 2:**
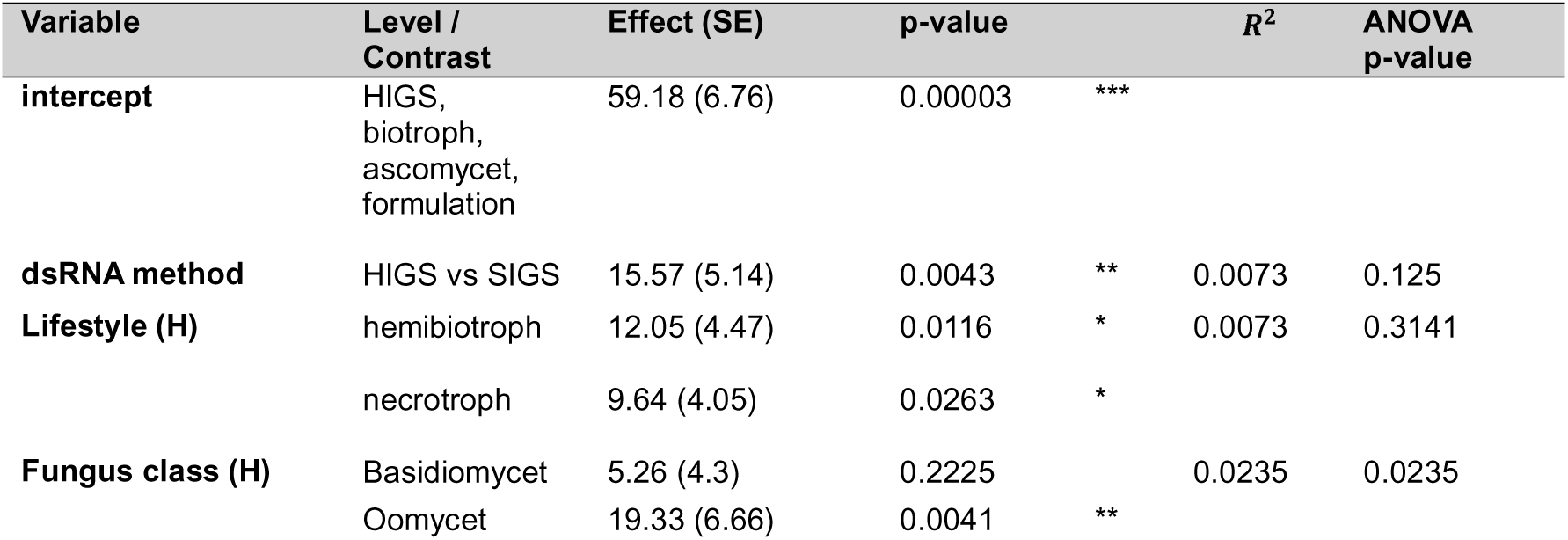
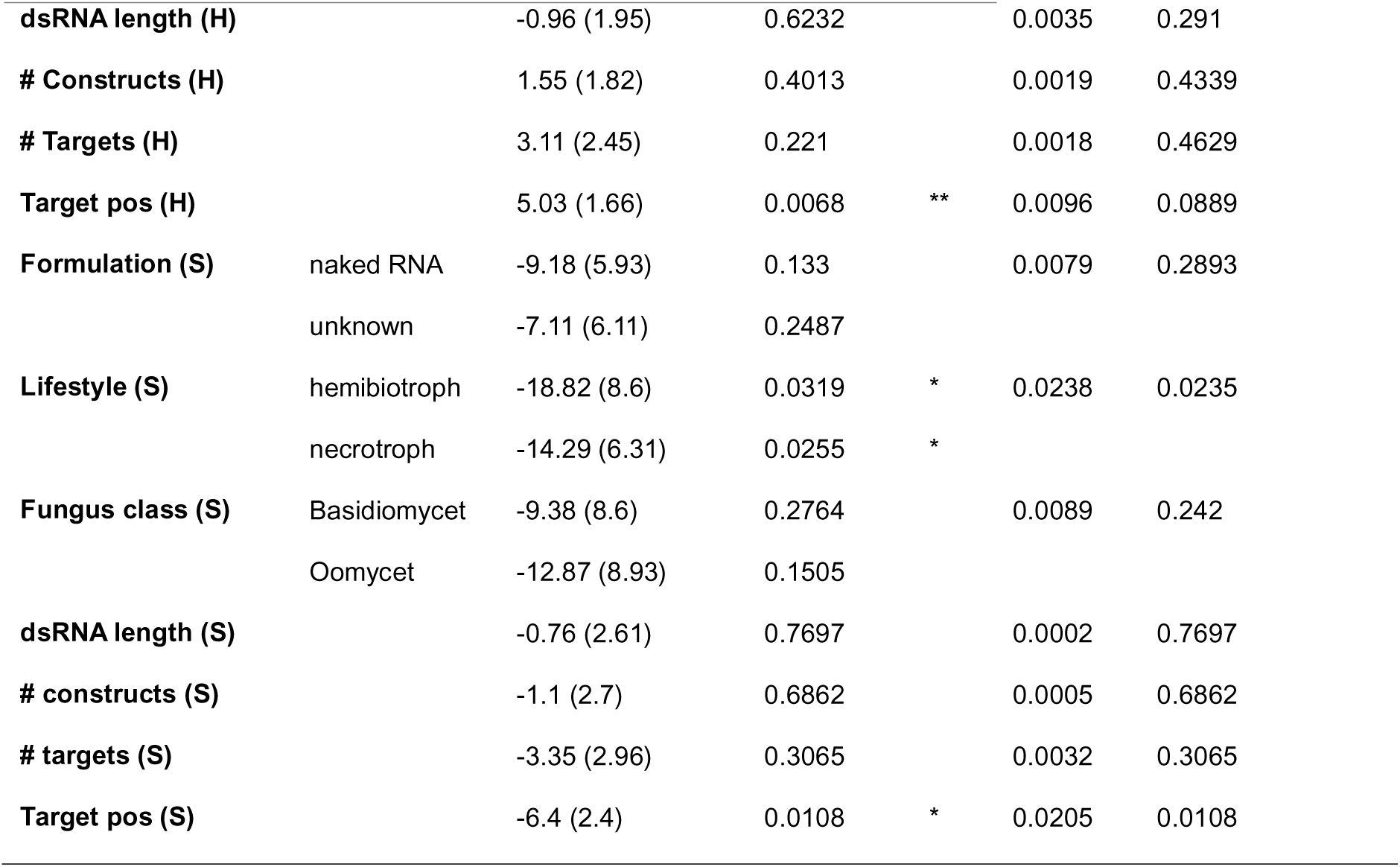
Results of the meta-regression model including type II ANOVA and relative contribution of fixed effects to the explained variance (R^2^). We separated R^2^ between fixed and random effects following (Nakagawa und Schielzeth 2013), with fixed effects (marginal R^2^) explaining approximately 12% of the variance in the data. Among all predictors, the fungal lifestyle in SIGS experiments contributed the most, accounting for 2.38% of the explained variance. Predictors are split by HIGS (H) and SIGS (S).

We believe the most likely reason for the lack of an effect when comparing between studies is the heterogeneity in experimental designs, including differences in plant species, pathogen strains, dsRNA concentrations, and application methods, which may mask the potential benefits of multiple targets. Additionally, many studies use RNAi design tools to preselect highly accessible and effective target sites; if individual constructs are already efficient, adding additional targets may not further enhance resistance. Finally, RNAi may have a threshold effect, where silencing a single essential gene is sufficient to achieve maximal resistance, limiting the observable impact of additional targets. Adding to this, using several constructs or a single construct with several targets while maintaining the same dsRNA concentration reduces the amount of dsRNA targeting each gene. This could hinder the resistance by reducing the dsRNA amount targeting an effective gene by adding targets that are potentially less effective.

However, combining several constructs may also provide additional benefits. For instance, combining multiple constructs can minimize off-target effects by using shorter and more selective dsRNA, while still achieving broad target coverage. Furthermore, combinatory dsRNAs can enhance specificity by targeting species-specific mRNA that, when applied together, produce effects comparable to those achieved by single conserved targets. This strategy may be relevant for pathogens with high genetic variability, as combining carefully selected targets increases the likelihood of effective silencing across different strains or isolates.

## Conclusions

Our meta-analysis of 89 published studies across diverse pathosystems revealed that SIGS confers overall higher resistance than HIGS. When fungal lifestyles were analyzed separately, SIGS was significantly more effective against biotrophic fungi, whereas HIGS conferred higher resistance in necrotrophic pathosystems. Although these trends were not always statistically significant for hemibiotrophs, likely because of limited data, the pattern was consistent across independent studies. Surprisingly, the use of dsRNA formulations in spray applications did not directly enhance initial resistance but rather prolonged its duration. Neither the length of the dsRNA nor the number of targeted genes had a significant impact on resistance outcomes. However, the position of the target site within the gene was relevant. Targeting near the 3′ end increased HIGS efficiency, whereas targeting near the 5′ end improved SIGS performance.

These findings suggest that RNA uptake mechanisms and infection biology are critical in shaping RNAi outcomes. In biotrophic interactions, haustorial contact between plants and fungi may facilitate the uptake of sprayed dsRNA in the apoplast or near stomata during early infection stages. High local dsRNA concentrations following spray application, combined with the intimate host association of biotrophic fungi, may enable efficient RNA uptake via pre-existing interfaces for transmembrane nutrient and effector exchange. Such uptake by germ tubes and haustorial precursors, likely promotes efficient RNA uptake and can trigger early gene silencing, disrupting infection establishment. Conversely, necrotrophic fungi destroy host tissue and feed on dead cells, making early dsRNA contact less critical. In this case, the continuous production of siRNAs by HIGS plants maintains a constitutive RNA supply that remains available during later infection phases, explaining the higher long-term resistance observed in necrotrophic systems.

These results provide the first quantitative evidence that SIGS outperforms HIGS against biotrophic fungi. They underscore the importance of adapting RNAi strategies to fungal lifestyles and indicate that target-specific optimization can further improve RNAi efficiency. However, further research on hemibiotrophic and transient biotrophic species, such as *Magnaporthe oryzae, Fusarium oxysporum,* and *Sclerotinia sclerotiorum* is needed to validate these patterns. The continuum of fungal lifestyles complicates classification and data interpretation, underlining the need for mechanistic studies on RNA uptake, processing, and cross-kingdom transfer. Our findings suggest that tailoring RNAi delivery to fungal lifestyles could significantly enhance RNAi-based crop protection. In particular, SIGS being non-transgenic, flexible, and effective against biotrophs, it represents a promising approach compatible with current regulatory frameworks. Advancing mechanistic understanding in fungi will be crucial for improving RNAi-based disease control and risk assessment, especially in the context of upcoming regulatory approval, such as the registration of the first RNA-based fungicide in Brazil.

## Authors contribution

PB, AS, FH and AK designed the analysis and conceptualized the project. PB, AS and JD acquired the data. PB, FH and AK analysed and interpreted the data. PB, JD, FH and AK wrote the manuscript. FH and AK were responsible for funding. All authors read and approved the manuscript.

## Data and code availability

The data that support the findings of this study are openly available in Zenodo at https://zenodo.org/records/17832226?token=eyJhbGciOiJIUzUxMiJ9.eyJpZCI6IjIxMWUzMjMzLWQwMzMtNDRmMi05NmUyLTI2ZTc0Y2M0MDAxNSIsImRhdGEiOnt9LCJyYW5kb20iOiIyMjM2YjVhMWM1OTY3NmQzZDBkYjZkZDk3MmQyN2Y0YiJ9.K96VipRCuN1nNTPOmP89lyEkMVaQ8xeXctNkGd0ZFe-PfeJ5TkypZCnDVCmZ_MUQjPnWQgU_eY171hR0hcfrJA, reference number Scripts used for data preparation, model fitting and plot creation are available at: https://github.com/TheoreticalEcology/RNAi-MetaAnalysis

## Supporting information

- Supporting Figures:

- Figure S1: Effect of number of constructs and targets on resistance
- Supporting Data:

- Data S1: Publications found by database query
- Data S2: Publications included in meta-analysis
- Data S3: Extracted figures and metaDigitize data
- Data S4: Extracted data
- Data S5: Processed data
- Supporting Methods:

- Method S1: Database queries
- Method S2: Inclusion criteria
- Method S3: Filtering of experiments
- Method S4: Data extraction
- Supporting Files:

- File S1: Data cleaning and imputation
- File S2: Data overview
- File S3: Meta-analysis and scatter plots

