## Supporting material for "Efficiency of RNAi based gene silencing in fungi - a review and meta-analysis"

### Supporting Figure S1: Effect plots of number of constructs and targets

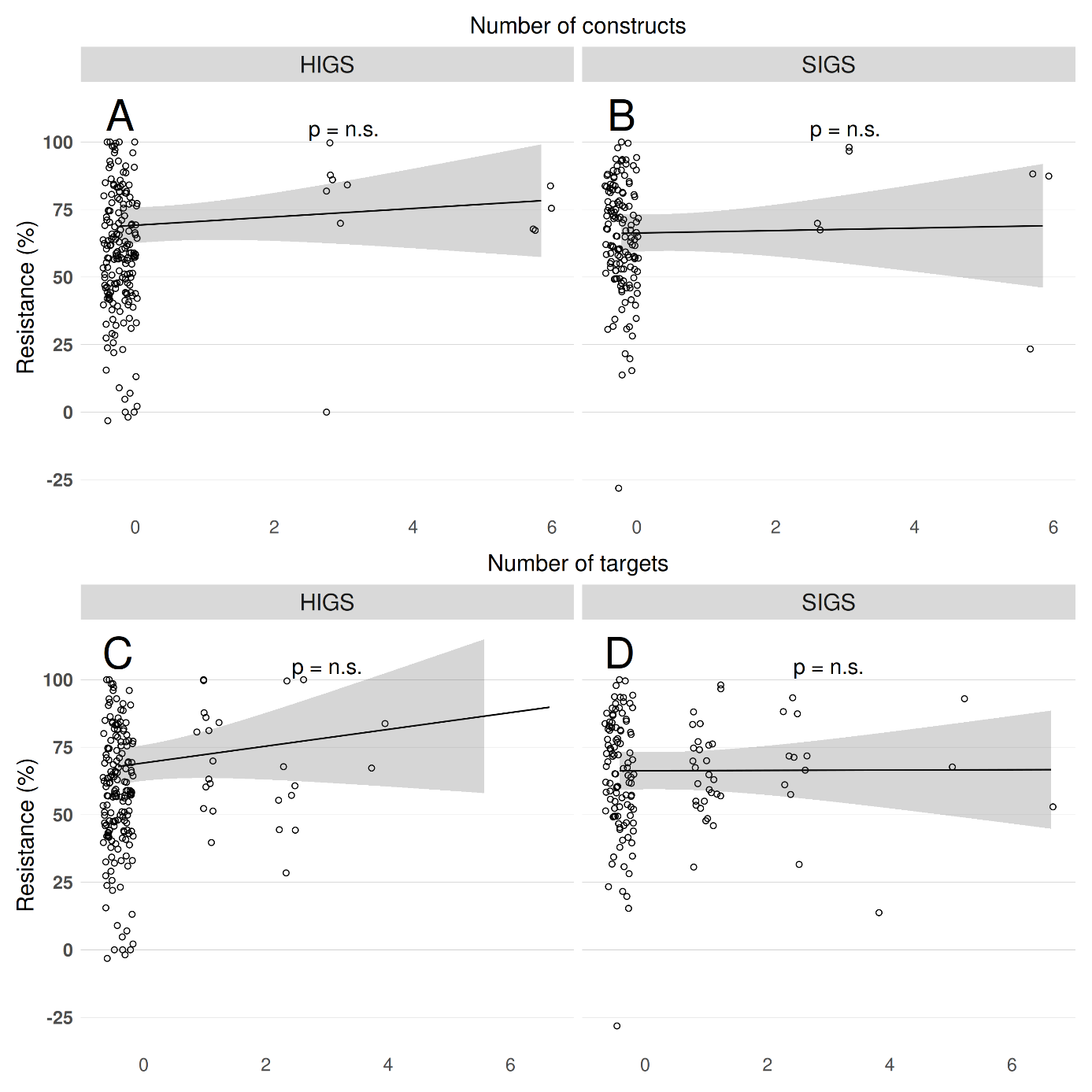
**Fig. S1 Effect plots of number of constructs and targets:** Effect plots showing the impact of the number of constructs in HIGS (A) and SIGS (B), as well as the number of targets in HIGS (C) and SIGS (D). Each point represents a distinct experiment. The solid line shows the predicted mean and the shaded area the 95% confidence band, both estimated using marginal means. Significance levels are based on the marginal means analysis: * P<0.05; ** P<0.01; *** P<0.001. Note that all displayed effects and p-values are marginal, i.e., the not adjusted for other covariates.

### Supporting Method S1: Database queries

To identify relevant publications for this meta-analysis, the PubMed database was queried.

The query term was designed to retrieve studies performing HIGS or SIGS experiments in which the plant resistance against fungi after RNAi treatment was measured. Review articles were excluded. The query was employed on August 5, 2024, with the following term and returned 71 publications (see supplementary file *Search_Results_05082024.csv*):

(((RNAi[Title/Abstract]) OR (siRNA[Title/Abstract]) OR (dsRNA[Title/Abstract])) AND (((plant[Title/Abstract]) OR (crop[Title/Abstract])) AND ((disease[Title/Abstract]) OR (pathogen*[Title/Abstract] OR (fung*[Title/Abstract])))) AND ((HIGS[Title/Abstract]) OR (Host-induced gene silencing[Title/Abstract]) OR (SIGS[Title/Abstract]) OR (Spray-induced gene silencing[Title/Abstract])) AND ((resistance[Title/Abstract]) OR (reduc*[Title/Abstract]) OR (inhibit*[Title/Abstract]) OR (efficien*[Title/Abstract]))) NOT (Review[Publication Type])

### Supporting Method S2: Inclusion criteria for fungal RNAi studies

#### Inclusion criteria for publications

In accordance with the Preferred Reporting Items for Systematic reviews and Meta-Analyses (PRISMA) guidelines (Page et al. 2021), the following inclusion and exclusion criteria were defined prior to the literature search:

1. The experiment was performed on fungi
2. The fungus was classified as necrotrophic, biotrophic, or hemibiotrophic
3. The experiment was qualified as either Host Induced or Spray Induced Gene Silencing (HIGS or SIGS)
4. Resistance was reported for both control and treatment groups, either as numerical or in a way from which a numerical can be derived
5. The target gene was a fungal gene
6. Actual resistance was reported
7. The experiments analyzed a fungus-host interaction
8. The fungal strain was not genetically modified
9. The plant cultivar used did not have specific resistances against the fungal strain
10. The resistance in at least one experiment was measured using at least one of the following criteria:
    a. Lesion size or affected leaf area (including pustule density/number and infection cushion area)
    b. Sporulation (including germination rate, number of sporangia/uredia and sclerotia formation)
    c. Fungal biomass (quantified via qPCR)
    d. Haustorium index or amount of haustoria (including appressorium and penetration rate)
    e. General disease reduction (including all measurements referring to disease severity, number of RFS/smut balls, number of affected leaves, or - in wheat - affected spikelets)

Consequently, the following exclusion criteria were applied:

1. Experiments that did not involve necrotrophic, biotrophic or hemibiotrophic fungi
2. Experiments employing methods other than HIGS or SIGS
3. Unreported control group and no additional information from which resistance could be deduced
4. Target gene was a plant gene
5. Only gene expression levels were reported, without resistance data
6. Experiments conducted only on fungal cultures in Petri dishes
7. Fungal strain was genetically modified
8. Experiments involved HIGS transgenic plants crossed with other cultivars

#### Definition of SIGS for data extraction

For the purpose of data extraction, SIGS was defined as the dsRNA application to the exterior of the plant. Consequently, experiments applying ssRNA (single stranded RNA) - in either sense, antisense, or both - as well as those involving the injection of dsRNA into plant tissue (e.g., via a syringe) were excluded.

#### Study selection

Each publication was independently assessed by a single reviewer. Initially, the abstract was screened to determine whether the publication met the predefined criteria. For instance, the abstract of Kamaraju et al mentions exclusively nematodes as RNAi targets, and was therefore excluded (Kamaraju et al. 2024).

If no decision could be made based on the abstract alone, the methods and results sections were examined to determine whether the publication contains at least one experiment suitable for the meta-analysis. If such experiments were identified, it was further assessed whether both a control as well as a treatment group were present, and whether numerical resistance values were either reported or could be deduced from the data.

The screening process led to the exclusion of 25 publications, resulting in 47 remaining studies. Details on all excluded publications are provided in supplementary file *Exclusion_List.xlsx*. In addition, 42 relevant publications were identified via manual online search and included in the meta-analysis, bringing the total number of studies to 89 (see supplementary file “*paper_list.xlsx*”).

### Supporting Method S3: Filtering of experiments

All selected publications were screened for experiments suitable for the meta-analysis, based on the criteria outlined in the supplementary section “Inclusion criteria”. To avoid overrepresentation of individual experiments, such as those repeated with only minor variations (e.g., different plant parts, time points, or promoters), an additional filtering step was performed. This filtering step is also aimed to exclude experiments testing marginal conditions or those testing a large number of target genes to identify suitable conditions or RNAi target candidates.

The following rules were applied as the final filtering step prior to data extraction:

1. Positive and negative controls were excluded.
2. If multiple plant lines were generated using the same construct, only the best-performing line was included. If several experiments were done with these plant lines, the overall best-performing line was selected.
3. If experiments were done with different plant generations, only the earliest available generation was used for data extraction.
4. If both in vivo and in vitro experiments were performed, only the in vivo data were used for data extraction.
5. If several time points (days past infection) were measured, only the time point with the highest observed resistance was used for data extraction. If another value summarizing all time points (e.g., AUDPC) was provided, it was preferred over the single time point.
6. If different assay types were performed (e.g., leaf assay, whole-plant assay, field assay), the assay providing the most realistic scenario was selected. Specifically, whole-plant assays were preferred over leaf assays, and field assays were preferred over laboratory conditions.
7. If multiple dsRNA designs or targets were tested initially, but only one or a few were used in subsequent experiments, only the latter were used for data extraction.
8. Experiments introducing additional adaptations intended to interfere with silencing were excluded.
9. Experiments using fungal strains that were not the primary target of the dsRNA were excluded.
10. Multiple distinct experiments performed on the same leaf were excluded.

An example of rule 2 can be found in He et al., who tested three different dsRNAs in HIGS experiments and generated two distinct transgenic plant lines per dsRNA (He et al. 2019). While line Fg00677 L1 showed better performance in overall symptom scores (40.67% vs. 34.73%; see Fig. 6B), line L2 performed considerably better in biomass measurements (31.67% vs. 63.87%; see Fig. 6C). Therefore, L2 was selected for data extraction. Similarly, line L6 was used for Fg08731-RNAi and line L4 for CYP51-RNAi. Wang et al. presents two generations of the same plant line in figure 1F (Wang et al. 2020). In accordance with rule 3, only generation T2 was used for data extraction, while T3 was omitted.

Xu et al. performed RNAi experiments using fungal strains that were not the intended target of the dsRNA (Xu et al. 2024); see Fig. 6I). Although significantly increased resistance against the fungus was observed, these experiments were excluded because the fungal strain was not the main target.

Pant et al. performed several distinct experiments on the same leaf (Pant und Kaur 2024); see Fig. 3). Since it cannot be ruled out that applied dsRNAs exert systemic effects that may influence neighboring experiments, all of these samples were excluded from data extraction.

### Supporting Method S4: Data extraction

#### Overview of extracted data

All experiments that fulfilled the inclusion criteria and did not violate any of the previously defined filtering rules were included for data extraction. When available, the following values and experimental details were extracted from the main text or the supplementary material of each publication:

1. Applied method (HIGS or SIGS).
2. Fungal classification: class, species, strain, and NCBI taxonomy ID.
3. Fungal lifestyle (biotroph, hemibiotroph, or necrotroph).
4. Plant classification: species, strain/cultivar, and NCBI taxonomy ID.
5. (HIGS only) Construct/plasmid and promoter used.
6. (HIGS only) Transformation method.
7. (SIGS only) Spray concentration (%).
8. (SIGS only) Formulation and formulation concentration (%).
9. Measured resistance (%), type of measurement, and standard error.
10. Source of resistance data (figure, table or text).
11. Time of measurement post-infection (dpi).
12. Targeted gene, gene ID, gene product, and CDS length.
13. Number of different dsRNAs used and number of different gene targets.
14. Length of dsRNA and target region within the gene.
15. Tool used for dsRNA design.
16. First author, year, title and DOI of the publication.

If resistance was measured using multiple methods under identical experimental conditions, all measurements were extracted. For example, Hu et al. measured resistance based on both pustule density and fungal biomass under the same conditions (Hu et al. 2020); see Fig. 4 and 6), and both values were included in the dataset.

In some cases, such as Haile et al., resistance was reported at multiple time points as well as in the form of an AUDPC (area under disease progress curve) (Haile et al. 2021); see Fig 1B). As the AUDPC integrates resistance over time into a single value, it was preferred over individual time point measurements for data extraction.

#### Resistance data extraction

The preferred method for resistance extraction was to identify direct numerical values reported in the text or tables. This also included cases in which plants did not show any symptoms after the treatment, such as in Dou et al. (Dou et al. 2020). These cases were recorded as 100% resistance.

If the same experimental results were presented in both figures and tables, only the numerical values from the tables were used, as extracting values from figures introduces minor uncertainties. For example, Pliego et al. also present all experimental results from table 1 in figure 1 (Pliego et al. 2013). Accordingly, only the table values were extracted. However, in cases where resistance values were reported numerically but without associated error values, the corresponding figures were used to extract both the resistance and the error value.

In some cases, standard errors were extremely small and could not be properly extracted, resulting in a value of 0. To address this, a small value of 0.001 was added to all standard errors before calculating model weights. The weights were calculated using the following formula:

$$weight= \frac{1}{{(standardError+0.001)}^{2}}$$

#### Calculation of resistance and error propagation

In most cases, resistance values were not reported directly. Instead, measurements were provided for both the control as well as the treated plants, which are then compared. Since resistance reflects the reduction of disease symptoms relative to the control, it was calculated using the following formula:

$$resistance \left( \% \right)=100-\left( \frac{{measurement}_{sample}}{{measurement}_{control}}*100 \right)$$

To account for uncertainty, standard error propagation was applied using the following formula:

$$SE= \frac{{measurement}_{sample}}{{measurement}_{control}}* \sqrt{\left( \frac{{SE}_{sample}}{{measurement}_{sample}} \right)^{2}+\left( \frac{{SE}_{control}}{{measurement}_{control}} \right)^{2}}$$

As many publications did not report measurements in numerical form but instead presented results via figures, data had to be extracted from these figures. For this purpose, the R package metaDigitize (Pick et al. 2019) was employed to extract the relevant values. To ensure reproducibility as well as traceability, all images used for data extraction, together with all corresponding data points, are provided in the supplementary dataset [add Zenodo link to data].

Lesion sizes were typically reported as area measurements. In some cases, they were reported as diameters, such as in Cheng et al., Table 1 (Cheng et al. 2015). To ensure comparability across studies, diameter values were converted to areas using the following formula:

$$area= \pi*\left( \frac{diameter}{2} \right)^{2}$$

An exception was made when the lesions exceeded the width of the leaf, as observed in Song et al., figure 7C (Song et al. 2018). In such cases, the original diameter value was retained.

McCaghey et al. reported measurements on a binary logarithmic scale (McCaghey et al. 2021). Therefore, all values were exponentiated with base 2 to recover the original measurement values.

#### Determination of the resistance value for the control

In cases where multiple control experiments were provided, the control condition that most closely matched the experimental conditions, while interfering as little as necessary, was chosen for resistance calculation. For example, Werner et al. provided control experiments using either TE buffer or GFP dsRNA (Werner et al. 2020). In this case, TE buffer was chosen as the control, as GFP dsRNA could potentially interfere with the experimental outcome. Similarly, Panwar et al. used both water inoculation and an empty plasmid as control experiments (Panwar et al. 2013); see Fig. 3). Since the empty plasmid more closely reflects the experimental conditions, it was chosen over water inoculation. Non-empty plasmids (e.g., expressing GFP dsRNA) were only considered as control experiments for resistance calculation when no other, less interfering control experiment was available.

#### Handling of missing uncertainties in the meta-analysis

When the variability or uncertainty of measurements or estimators (e.g., standard error or standard deviation of the group mean) was not explicitly stated for a given figure, it was examined whether the publication specified these values for a similar figure. If so, it was assumed that experimental variability was consistent across figures. For example, Sharma et al. did not specify a metric of variance or uncertainty in Fig. 3, but stated standard error in Fig. 2 (Sharma et al. 2018). Therefore, standard error was also assumed for figure 3 to maintain consistency.

In cases where no information was available, we used imputed values for the standard error (e.g. in Zhang et al. 2019, or in Fig. 4E in Xu et al. 2019).

All of such occurrences were documented in the data extraction R script (*extract_data_from_figures.R*).

#### Target gene and dsRNA information

If no gene ID was provided for the RNAi target NCBI BLASTn (Camacho et al. 2009) was used with all available information, such as the dsRNA sequence (if provided) or the primer sequences used to generate the dsRNAs. If no suitable match could be identified via BLASTn, additional databases such as MycoCosm (Nordberg et al. 2014) or Ensembl (Dyer et al. 2025) were used with any available information. If a target could be identified, the associated nucleotide sequence was used to determine:

- The target gene length
- The targeted position within the gene
- The dsRNA length (if not explicitly mentioned in the manuscript)

For the latter, the positions of the forward and reverse primers on the target sequence were used to calculate the dsRNA length. If possible, only the CDS length of the target gene was extracted.

#### Calculate relative coverage

The relative coverage of the dsRNA on the target gene was calculated using the following formula:

$$relativeCoverage \left( \% \right)=100* \left( \frac{dsRNAlength}{targetGeneLength} \right)$$

#### Calculate relative position of dsRNA on target gene

The relative mean position of the dsRNA within the target gene(s) was calculated using the following formula:

$$mean position \left( \% \right) = \frac{\left( \frac{startPosition + endPosition}{2} \right)}{lengthTargetGene}\cdot100$$
