## Supporting files and data for "Efficiency of RNAi based gene silencing in fungi - a review and meta-analysis": 01-data_editing.html

Meta-analyse HIGS vs SIGS


Code 

- Show All Code
- Hide All Code

### Meta-analyse HIGS vs SIGS

###### Patrick Barth

Abstract

This markdown documents prepares data gathered from publications which
examined the applied resistance to plants against fungi via HIGS or SIGS
experiments. The preparation involves calculating the mean for entries
with several values, the calculation of the mean position of the dsRNA
upon the target mRNA, the merging of experiments that differ only in the
dsRNA concentration used (only SIGS) or the measurement method and the
imputation of missing data points.

### Set up

#### Set seed

```
set.seed(123)
```

#### Load libraries

```
library(stringr)
library(ggpubr)
```

```
## Loading required package: ggplot2
```

```
library(missRanger) # Imputation (fill in NAs in data sets)
```

#### Functions

##### Calculate mean value

Function to calculate the mean value of numerical entries split by a
semicolon

```
get_mean_for_all_with_semicolon <-function(df, var) {
  rows <- which(grepl(';',df[,var]))
  df[rows,var] <- unlist(lapply(df[rows,var], function(x){
    values <- split_numerical_entries(x)
    mean_value <- round(mean(values))
    return(mean_value)
  }))
  return(df)
}
```

##### Split numerical entries

Function to split entries separated by a semicolon

```
split_numerical_entries <- function(x) {
  values <- unlist(strsplit(x,';'))
  values <- as.numeric(values[!is.na(values)])
  return(values)
}
```

##### Compare all rows of current column

Function that compares all entries of a column with another value and
returns TRUE if the value is the same

```
compare_rows <- function(df,row,column) {
  return(df[,column] == df[row,column] 
         & !is.na(df[,column])
         &!is.na(df[row,column])
         | (is.na(df[,column]) & is.na(df[row,column])))
}
```

##### Check merged entries

Function to check all entries of a column contain the same value as
the first row. Writes a merged statement if they differ

```
check_merged_entries <- function(df,column) {
  if (all(df[1,column] == df[,column], na.rm = TRUE)
      & !any(is.na(df[, column]))) {
    return(df[1,column])
  } else {
    return('sample got merged. see raw data')
  }
}
```

#### Read data

Read raw data

```
raw_table <- read.table(file = "../data/data_current.csv",
                        header = TRUE,
                        fill = TRUE,
                        sep='\t')
```

#### Prepare data

##### Calculate mean values of single entries

Calculate the mean for numerical entries containing several
values.

```
edit_table <- raw_table

# Calculate mean for all samples with multiple entries
edit_table <- get_mean_for_all_with_semicolon(edit_table,"dsRNA_length")
edit_table <- get_mean_for_all_with_semicolon(edit_table,"target_region_start")
```

```
## Warning in split_numerical_entries(x): NAs introduced by coercion
## Warning in split_numerical_entries(x): NAs introduced by coercion
## Warning in split_numerical_entries(x): NAs introduced by coercion
## Warning in split_numerical_entries(x): NAs introduced by coercion
```

```
edit_table <- get_mean_for_all_with_semicolon(edit_table,"target_region_end")
```

```
## Warning in split_numerical_entries(x): NAs introduced by coercion
## Warning in split_numerical_entries(x): NAs introduced by coercion
## Warning in split_numerical_entries(x): NAs introduced by coercion
## Warning in split_numerical_entries(x): NAs introduced by coercion
```

```
edit_table <- get_mean_for_all_with_semicolon(edit_table,"target_length")
```

```
## Warning in split_numerical_entries(x): NAs introduced by coercion
## Warning in split_numerical_entries(x): NAs introduced by coercion
## Warning in split_numerical_entries(x): NAs introduced by coercion
```

##### Calculate mean position

Calculates mean position (in %) of the dsRNA upon the target mRNA by
including the start and end position as well as the length of the target
mRNA.

```
edit_table$target_mean_percent <- apply(edit_table,1, function(x){
  start_pos     <- x['target_region_start']
  end_pos       <- x['target_region_end']
  target_length <- x['target_length']
  
# Calculate mean position in % of dsRNA position on target mRNA
if(is.na(start_pos) | is.na(end_pos) | is.na(target_length)){
  result <- NA
} else {
  mean <- round((as.double(start_pos) + as.double(end_pos))/2)
  result <- round((mean/as.double(target_length))*100)
}
  return(result)
})
```

```
## Warning in FUN(newX[, i], ...): NAs introduced by coercion
## Warning in FUN(newX[, i], ...): NAs introduced by coercion
## Warning in FUN(newX[, i], ...): NAs introduced by coercion
## Warning in FUN(newX[, i], ...): NAs introduced by coercion
## Warning in FUN(newX[, i], ...): NAs introduced by coercion
## Warning in FUN(newX[, i], ...): NAs introduced by coercion
```

##### Remove emtpy entries

Remove entries that provide no resistance measurement

```
# Remove potential entries without resistance
edit_table <- edit_table <- edit_table[!is.na(edit_table$resistance) & trimws(edit_table$resistance) != "", ]
```

##### Set correct types for entries

Set type factor for nominal and double for numerical entries

```
edit_table$dsRNA_method <- as.factor(edit_table$dsRNA_method)
edit_table$dsRNA_length <- as.double(edit_table$dsRNA_length)
edit_table$lifestyle <- as.factor(edit_table$lifestyle)
edit_table$number_of_constructs <- as.factor(edit_table$number_of_constructs)
edit_table$number_of_targets <- as.factor(edit_table$number_of_targets)
edit_table$fungus_class <- as.factor(edit_table$fungus_class)
edit_table$formulation <- as.factor(edit_table$formulation)
edit_table$target_region_start <- as.double(edit_table$target_region_start)
```

```
## Warning: NAs introduced by coercion
```

```
edit_table$target_region_end <- as.double(edit_table$target_region_end)
```

```
## Warning: NAs introduced by coercion
```

```
edit_table$target_length <- as.double(edit_table$target_length)
edit_table$prediction_tool <- as.factor(edit_table$prediction_tool)
edit_table$standard_error <- as.double(edit_table$standard_error)
```

```
## Warning: NAs introduced by coercion
```

```
edit_table$resistance <- as.double(edit_table$resistance)
```

##### Summarize extraction methods

Summarize different methods of resistance measurement in order to
reduce them to: General reduction, Infected leaf area, Biomass,
Haustoria and Sporulation

```
# Merge resistance extraction method based on their overall group
edit_table[which(edit_table$resistance_measurement == "disease reduction" |
                 edit_table$resistance_measurement == "reduction" |
                 edit_table$resistance_measurement == "number smut balls" |
                 edit_table$resistance_measurement == "Number RFS balls" |
                 edit_table$resistance_measurement == "number diseased leaves" |
                 edit_table$resistance_measurement == "number infected spikelets" |
                 edit_table$resistance_measurement == "number yellow leaves" |
                 edit_table$resistance_measurement == "number leaves with symptoms"),"measurement_combined"] <- "General reduction"

edit_table[which(edit_table$resistance_measurement == "lesion area" |
                 edit_table$resistance_measurement == "lesion density" |
                 edit_table$resistance_measurement == "infection area" |
                 edit_table$resistance_measurement == "leaf area" |
                 edit_table$resistance_measurement == "lesion length" |
                 edit_table$resistance_measurement == "pustule density" |
                 edit_table$resistance_measurement == "Infection cushion area"),"measurement_combined"] <- "Infected leaf area"

edit_table[which(edit_table$resistance_measurement == "biomass" ),"measurement_combined"] <- "Biomass"

edit_table[which(edit_table$resistance_measurement == "Appressorium" |
                 edit_table$resistance_measurement == "haustoria" |
                 edit_table$resistance_measurement == "penetration rate" ),"measurement_combined"] <- "Haustoria"

edit_table[which(edit_table$resistance_measurement == "germination rate" |
                 edit_table$resistance_measurement == "sclerotia formation" |
                 edit_table$resistance_measurement == "number of uredia" |
                 edit_table$resistance_measurement == "sporulation" ),"measurement_combined"] <- "Sporulation"
```

##### Summarize formulation

In some SIGS experiments a formulation is used in order to either
protect the dsRNA from environmental influences or to enable an easier
entry into the plants. As multiple different formulations were used, the
sample size for each formulation is too small to reach solid statistical
conclusions. Therefore, all usages of a formulation are summarized into
the group ‘formulation’. Furthermore, all SIGS experiments using naked
dsRNA are summarized into the group naked dsRNA and all SIGS experiments
not stating whether a formulation was used or not are summarized into
the group ‘unknown’.

```
# unify formulations
edit_table$formulation <- as.character(edit_table$formulation)
edit_table$formulation_merged <- ifelse(!is.na(edit_table$formulation) & 
                                      (edit_table$formulation == 'naked dsRNA' | 
                                       edit_table$formulation == 'naked' | 
                                       edit_table$formulation == 'Silwet L-77' | 
                                       edit_table$formulation == 'tween 20'), 
                                      'naked_dsRNA', 
                                      NA)

edit_table$formulation_merged <- ifelse(!is.na(edit_table$formulation) & 
                                      (edit_table$formulation == 'LDH' 
                    | edit_table$formulation == 'Nanoparticle'
                    | edit_table$formulation == 'artificial vesicle'
                    | edit_table$formulation == 'BioClay'
                    | edit_table$formulation == 'Diethylpyrocarbonate'
                    | edit_table$formulation == 'Nanoclay particles')
                    & (edit_table$formulation != 'naked'
                       & edit_table$formulation != 'naked dsRNA'), 
                                      'formulation', 
                                      edit_table$formulation_merged)

edit_table$formulation_merged <- ifelse(edit_table$dsRNA_method == 'SIGS' &
                                      ( is.na(edit_table$formulation)
                                      | edit_table$formulation == 'n.a.'
                                      | edit_table$formulation == 'n.a'
                                      | edit_table$formulation == ''), 
                                      'unknown', 
                                      edit_table$formulation_merged)

edit_table$formulation_merged <- as.factor(edit_table$formulation_merged)
```

##### Merge entries with differing concentrations

As some publications perform the same (SIGS) experiments with
different spray concentrations. These would introduce biases towards
such studies as well as keep studies that use very low concentrations on
purpose in order to analyse how low of a concentration is still
effective. Therefore, all experiments are compared based a number of
data points in order to determine all experiments which only differ in
the applied dsRNA concentration. These experiments are merged into a
single representative entry using the highest applied resistance of all
with the corresponding standard error.

```
edit_table$row <- 1:nrow(edit_table)
# Get all entries that are equal in a vast amount of predictors but different in either spray or formulation concentration
affected_entries <- data.frame(matrix(ncol = ncol(edit_table) , nrow = 0))
for (row in 1:nrow(edit_table)) {
  entries_to_extract <- which(edit_table$dsRNA_method == 'SIGS'
                            & compare_rows(edit_table,row,'fungus_species') 
                            & compare_rows(edit_table,row,'fungus_strain') 
                            & compare_rows(edit_table,row,'host_species') 
                            & compare_rows(edit_table,row,'host_strain') 
                            & compare_rows(edit_table,row,'target_gene') 
                            & compare_rows(edit_table,row,'dsRNA_length')
                            & compare_rows(edit_table,row,'number_of_constructs')
                            & compare_rows(edit_table,row,'number_of_targets')
                            & compare_rows(edit_table,row,'target_region_start')
                            & compare_rows(edit_table,row,'target_region_end')
                            & compare_rows(edit_table,row,'gene_ID')
                            & compare_rows(edit_table,row,'Assay')
                            & compare_rows(edit_table,row,'measurement_combined')
                            & compare_rows(edit_table,row,'formulation')
                            & edit_table[,'study_number'] == edit_table[row,"study_number"]
                            & ((edit_table[,'formulation_concentration'] != edit_table[row,"formulation_concentration"]
                                | edit_table[,'spray_concentration.ng.µl.'] != edit_table[row,"spray_concentration.ng.µl."])
                              & (!is.na(edit_table[,'formulation_concentration']) | !is.na(edit_table[row,"spray_concentration.ng.µl."]))
                              & (edit_table[,'formulation_concentration'] != '' | edit_table[row,"spray_concentration.ng.µl."] != ''))
                            & 1:nrow(edit_table) != row)
  other_entries <- paste(sort(c(row, entries_to_extract)), collapse = ';')
  new_entries <- edit_table[entries_to_extract,]
  if (nrow(new_entries) > 0) {
    new_entries$dup_with <- other_entries
    affected_entries <- rbind(affected_entries, new_entries)
  }
}

affected_entries <- unique(affected_entries)
for (current_entries in unique(affected_entries$dup_with)) {
  current_rows <- split_numerical_entries(current_entries)
  current_entries <- edit_table[which(edit_table$row %in% current_rows),]
  merged_entry <- current_entries[1,]
  
  merged_entry$resistance <- current_entries[which.max(current_entries$resistance),"resistance"]
  merged_entry$standard_error <- current_entries[which.max(current_entries$resistance),"standard_error"]
  
  merged_entry$formulation_concentration <- check_merged_entries(current_entries,"formulation_concentration")
  merged_entry$spray_concentration.ng.µl. <- check_merged_entries(current_entries,"spray_concentration.ng.µl.")
  merged_entry$sample <- check_merged_entries(current_entries,"sample")
  merged_entry$figure <- check_merged_entries(current_entries,"figure")

  # Entferne alte Eintrage und fuege den neuen gemergten hinzu
  edit_table <- edit_table[which(!edit_table$row %in% current_rows) ,]
  edit_table <- rbind(edit_table, merged_entry)
}
```

##### Merge entries that are basically the same experiment but with different measurement methods

Similar to the previous step some experiments are performed in the
same way only differing in the method used to measure the applied
resistance. These would introduce biases towards studies performing
multiple measurement methods. Therefore, all experiments are compared on
a number of data points in order to determine all experiments which only
differ in the applied measurement method. These experiments are merged
into a single representative entry using the mean value of the applied
resistance and standard error.

```
edit_table$row <- 1:nrow(edit_table)
# Get all entries that are equal in a vast amount of predictors but different in either spray or formulation concentration
affected_entries <- data.frame(matrix(ncol = ncol(edit_table) , nrow = 0))
for (row in 1:nrow(edit_table)) {
  entries_to_extract <- which(compare_rows(edit_table,row,'dsRNA_method') 
                            & compare_rows(edit_table,row,'fungus_species') 
                            & compare_rows(edit_table,row,'fungus_strain') 
                            & compare_rows(edit_table,row,'host_species') 
                            & compare_rows(edit_table,row,'host_strain') 
                            & compare_rows(edit_table,row,'target_gene') 
                            & compare_rows(edit_table,row,'dsRNA_length')
                            & compare_rows(edit_table,row,'number_of_constructs')
                            & compare_rows(edit_table,row,'number_of_targets')
                            & compare_rows(edit_table,row,'target_region_start')
                            & compare_rows(edit_table,row,'target_region_end')
                            & compare_rows(edit_table,row,'gene_ID')
                            & compare_rows(edit_table,row,'Assay')
                            & compare_rows(edit_table,row,'formulation')
                            & compare_rows(edit_table,row,'formulation_concentration')
                            & compare_rows(edit_table,row,'spray_concentration.ng.µl.')
                            & edit_table[,'study_number'] == edit_table[row,"study_number"]
                            & 1:nrow(edit_table) != row)
  other_entries <- paste(sort(c(row, entries_to_extract)), collapse = ';')
  new_entries <- edit_table[entries_to_extract,]
  if (nrow(new_entries) > 0) {
    new_entries$dup_with <- other_entries
    affected_entries <- rbind(affected_entries, new_entries)
  }
}

affected_entries <- unique(affected_entries)
for (current_entries in unique(affected_entries$dup_with)) {
  current_rows <- split_numerical_entries(current_entries)
  current_entries <- edit_table[which(edit_table$row %in% current_rows),]
  merged_entry <- current_entries[1,]
  
  merged_entry$resistance <- mean(current_entries$resistance)
  merged_entry$standard_error <- mean(current_entries$standard_error)
  
  merged_entry$resistance_measurement <- 'sample got merged. see raw data'
  
  merged_entry$construct.plasmid <- check_merged_entries(current_entries,"construct.plasmid")
  merged_entry$promotor <- check_merged_entries(current_entries,"promotor")
  merged_entry$transformation_method <- check_merged_entries(current_entries,"transformation_method")
  merged_entry$sample <- check_merged_entries(current_entries,"sample")
  merged_entry$figure <- check_merged_entries(current_entries,"figure")

  # Entferne alte Eintrage und fuege den neuen gemergten hinzu
  edit_table <- edit_table[which(!edit_table$row %in% current_rows) ,]
  edit_table <- rbind(edit_table, merged_entry)
}
edit_table$row <- 1:nrow(edit_table)
```

##### data imputation

Determine the amount of entries that contain at least one NA in one
of the columns on which imputation is performed

```
print("Number of entries containing at least one NA in one of the predictors:")
```

```
## [1] "Number of entries containing at least one NA in one of the predictors:"
```

```
sum(rowSums(is.na(edit_table[,c("standard_error","dsRNA_length","formulation_merged","target_length","target_mean_percent","number_of_constructs","number_of_targets")])) > 0)
```

```
## [1] 240
```

```
print("Total number of NAs present in all predictors (out of a total of 2401 values):")
```

```
## [1] "Total number of NAs present in all predictors (out of a total of 2401 values):"
```

```
sum(rowSums(is.na(edit_table[,c("standard_error","dsRNA_length","formulation_merged","target_length","target_mean_percent","number_of_constructs","number_of_targets")])))
```

```
## [1] 436
```

Since a multitude of experiments are missing single or multiple data
points data imputation is performed in order to guarantee a correct data
basis for the subsequent linear model. Therefore, data imputation is
performed for all predictors used in the later model as well as for the
standard error which was not supplied in all experiments.

```
edit_table$id <- 1:nrow(edit_table)

edit_table[,c("standard_error_imputed","dsRNA_length_imputed","formulation_merged_imputed","target_length_imputed","target_mean_percent_imputed","number_of_constructs_imputed","number_of_targets_imputed")] <- edit_table[,c("standard_error","dsRNA_length","formulation_merged","target_length","target_mean_percent","number_of_constructs","number_of_targets")]


# impute missing data
edit_table <- missRanger(edit_table,
                         formula =  standard_error_imputed + dsRNA_length_imputed + formulation_merged_imputed 
                            + target_length_imputed + target_mean_percent_imputed 
                            + number_of_constructs_imputed + number_of_targets_imputed 
                         ~ resistance + dsRNA_method + fungus_class + lifestyle + standard_error_imputed 
                          + dsRNA_length_imputed + formulation_merged_imputed + target_length_imputed 
                          + target_mean_percent_imputed + number_of_constructs_imputed + number_of_targets_imputed)
```

```
## 
## Variables to impute:     number_of_constructs_imputed, number_of_targets_imputed, dsRNA_length_imputed, target_length_imputed, standard_error_imputed, target_mean_percent_imputed, formulation_merged_imputed
## Variables used to impute:    resistance, dsRNA_method, fungus_class, lifestyle, standard_error_imputed, dsRNA_length_imputed, formulation_merged_imputed, target_length_imputed, target_mean_percent_imputed, number_of_constructs_imputed, number_of_targets_imputed
## 
## iter 1 
##   |                                                                              |                                                                      |   0%  |                                                                              |==========                                                            |  14%  |                                                                              |====================                                                  |  29%  |                                                                              |==============================                                        |  43%  |                                                                              |========================================                              |  57%  |                                                                              |==================================================                    |  71%  |                                                                              |============================================================          |  86%  |                                                                              |======================================================================| 100%
## iter 2 
##   |                                                                              |                                                                      |   0%  |                                                                              |==========                                                            |  14%  |                                                                              |====================                                                  |  29%  |                                                                              |==============================                                        |  43%  |                                                                              |========================================                              |  57%  |                                                                              |==================================================                    |  71%  |                                                                              |============================================================          |  86%  |                                                                              |======================================================================| 100%
## iter 3 
##   |                                                                              |                                                                      |   0%  |                                                                              |==========                                                            |  14%  |                                                                              |====================                                                  |  29%  |                                                                              |==============================                                        |  43%  |                                                                              |========================================                              |  57%  |                                                                              |==================================================                    |  71%  |                                                                              |============================================================          |  86%  |                                                                              |======================================================================| 100%
## iter 4 
##   |                                                                              |                                                                      |   0%  |                                                                              |==========                                                            |  14%  |                                                                              |====================                                                  |  29%  |                                                                              |==============================                                        |  43%  |                                                                              |========================================                              |  57%  |                                                                              |==================================================                    |  71%  |                                                                              |============================================================          |  86%  |                                                                              |======================================================================| 100%
```

remove some unnecessary columns

```
edit_table <- edit_table[,-which(names(edit_table) %in% c("comment","comment2","comment3","problems","problems2","row"))]
```

##### Export data

Set correct data types

```
edit_table$number_of_constructs <- as.integer(edit_table$number_of_constructs)
edit_table$number_of_targets <- as.integer(edit_table$number_of_targets)
edit_table$formulation <- as.factor(edit_table$formulation)
edit_table$number_of_constructs_imputed <- as.double(edit_table$number_of_constructs_imputed)
edit_table$number_of_targets_imputed <- as.double(edit_table$number_of_targets_imputed)
```

Scale numerical predictors

```
edit_table$sDsRNA_length <- scale(edit_table$dsRNA_length_imputed)
edit_table$sNumber_of_constructs <- scale(edit_table$number_of_constructs_imputed)
edit_table$sNumber_of_targets <- scale(edit_table$number_of_targets_imputed)
edit_table$sTarget_mean <- scale(edit_table$target_mean_percent_imputed)
```

Rename predictors used in the model

```
names(edit_table)[names(edit_table) == "dsRNA_method"] <- "fDsRNA_method"
names(edit_table)[names(edit_table) == "lifestyle"] <- "fLifestyle"
names(edit_table)[names(edit_table) == "fungus_class"] <- "fFungus_class"
names(edit_table)[names(edit_table) == "formulation_merged_imputed"] <- "fFormulation"
```

```
save(edit_table, file = "../data/data_edited.RData")
write.table(edit_table,
            file = "../data/data_edited.csv",
            sep = "\t",
            row.names = FALSE)
```

```
Sys.Date()
## [1] "2025-12-08"
sessionInfo()
## R version 4.3.3 (2024-02-29)
## Platform: x86_64-pc-linux-gnu (64-bit)
## Running under: Ubuntu 24.04.3 LTS
## 
## Matrix products: default
## BLAS:   /usr/lib/x86_64-linux-gnu/blas/libblas.so.3.12.0 
## LAPACK: /usr/lib/x86_64-linux-gnu/lapack/liblapack.so.3.12.0
## 
## locale:
##  [1] LC_CTYPE=en_US.UTF-8       LC_NUMERIC=C              
##  [3] LC_TIME=en_US.UTF-8        LC_COLLATE=en_US.UTF-8    
##  [5] LC_MONETARY=en_US.UTF-8    LC_MESSAGES=en_US.UTF-8   
##  [7] LC_PAPER=en_US.UTF-8       LC_NAME=C                 
##  [9] LC_ADDRESS=C               LC_TELEPHONE=C            
## [11] LC_MEASUREMENT=en_US.UTF-8 LC_IDENTIFICATION=C       
## 
## time zone: Europe/Berlin
## tzcode source: system (glibc)
## 
## attached base packages:
## [1] stats     graphics  grDevices utils     datasets  methods   base     
## 
## other attached packages:
## [1] missRanger_2.6.1 ggpubr_0.6.0     ggplot2_3.5.2    stringr_1.5.1   
## 
## loaded via a namespace (and not attached):
##  [1] Matrix_1.6-5       gtable_0.3.6       jsonlite_2.0.0     ranger_0.17.0     
##  [5] dplyr_1.1.4        compiler_4.3.3     ggsignif_0.6.4     Rcpp_1.0.14       
##  [9] tidyselect_1.2.1   tidyr_1.3.1        jquerylib_0.1.4    scales_1.4.0      
## [13] yaml_2.3.10        fastmap_1.2.0      lattice_0.22-5     R6_2.6.1          
## [17] generics_0.1.4     Formula_1.2-5      knitr_1.50         backports_1.5.0   
## [21] tibble_3.2.1       car_3.1-3          bslib_0.9.0        pillar_1.10.2     
## [25] RColorBrewer_1.1-3 rlang_1.1.6        cachem_1.1.0       broom_1.0.8       
## [29] stringi_1.8.7      xfun_0.52          sass_0.4.10        cli_3.6.5         
## [33] withr_3.0.2        magrittr_2.0.3     digest_0.6.37      grid_4.3.3        
## [37] rstudioapi_0.17.1  lifecycle_1.0.4    vctrs_0.6.5        rstatix_0.7.2     
## [41] evaluate_1.0.3     glue_1.8.0         farver_2.1.2       abind_1.4-8       
## [45] carData_3.0-5      rmarkdown_2.29     purrr_1.0.4        tools_4.3.3       
## [49] pkgconfig_2.0.3    htmltools_0.5.8.1
```
