## Supporting files and data for "Efficiency of RNAi based gene silencing in fungi - a review and meta-analysis": 02-data_overview.html

Meta-analyse HIGS vs SIGS


Code 

- Show All Code
- Hide All Code

### Meta-analyse HIGS vs SIGS

###### Patrick Barth

Abstract

This Markdown document generates an overview of the sample sizes and
means of different data splits.

### Set up

#### overview of observations

Amount of entries split by dsRNA method and fungal lifestyle

```
##               
##                HIGS SIGS Sum
##   biotroph       75   58 133
##   hemibiotroph   48   10  58
##   necrotroph     72   80 152
##   Sum           195  148 343
```

Mean resistance and standard error of resistance measurements split
by dsRNA method and fungal lifestyle

```
##       lifestyle method     mean       SE
## 1      biotroph   HIGS 55.20272 2.450545
## 2      biotroph   SIGS 70.26440 2.607690
## 3      biotroph    Sum 61.77097 1.897783
## 4  hemibiotroph   HIGS 65.08280 4.177184
## 5  hemibiotroph   SIGS 63.93047 2.672416
## 6  hemibiotroph    Sum 64.88412 3.479192
## 7    necrotroph   HIGS 59.98482 2.255183
## 8    necrotroph   SIGS 61.49388 2.424996
## 9    necrotroph    Sum 60.77906 1.660053
## 10          all   HIGS 59.40044 1.638715
## 11          all   SIGS 65.09561 1.700206
## 12          all    Sum 61.85783 1.193899
```

### Reproducibility info

```
Sys.Date()
## [1] "2025-12-08"
sessionInfo()
## R version 4.3.3 (2024-02-29)
## Platform: x86_64-pc-linux-gnu (64-bit)
## Running under: Ubuntu 24.04.3 LTS
## 
## Matrix products: default
## BLAS:   /usr/lib/x86_64-linux-gnu/blas/libblas.so.3.12.0 
## LAPACK: /usr/lib/x86_64-linux-gnu/lapack/liblapack.so.3.12.0
## 
## locale:
##  [1] LC_CTYPE=en_US.UTF-8       LC_NUMERIC=C              
##  [3] LC_TIME=en_US.UTF-8        LC_COLLATE=en_US.UTF-8    
##  [5] LC_MONETARY=en_US.UTF-8    LC_MESSAGES=en_US.UTF-8   
##  [7] LC_PAPER=en_US.UTF-8       LC_NAME=C                 
##  [9] LC_ADDRESS=C               LC_TELEPHONE=C            
## [11] LC_MEASUREMENT=en_US.UTF-8 LC_IDENTIFICATION=C       
## 
## time zone: Europe/Berlin
## tzcode source: system (glibc)
## 
## attached base packages:
## [1] stats     graphics  grDevices utils     datasets  methods   base     
## 
## loaded via a namespace (and not attached):
##  [1] digest_0.6.37     R6_2.6.1          fastmap_1.2.0     xfun_0.52        
##  [5] cachem_1.1.0      knitr_1.50        htmltools_0.5.8.1 rmarkdown_2.29   
##  [9] lifecycle_1.0.4   cli_3.6.5         sass_0.4.10       jquerylib_0.1.4  
## [13] compiler_4.3.3    rstudioapi_0.17.1 tools_4.3.3       evaluate_1.0.3   
## [17] bslib_0.9.0       yaml_2.3.10       rlang_1.1.6       jsonlite_2.0.0
```
