## Supporting files and data for "Efficiency of RNAi based gene silencing in fungi - a review and meta-analysis": 02-data_overview.Rmd

````
---
author: 'Patrick Barth'
title: 'Meta-analyse HIGS vs SIGS'
abstract: This Markdown document generates an overview of the sample sizes and means of different data splits.
output: 
  html_document:
    code_folding: 'show'
editor_options: 
  chunk_output_type: console
---


# Set up


```{r session info,collapse=TRUE, echo=FALSE}
set.seed(123)
```

```{r, echo=FALSE}
load("../data/data_edited.RData")
```


```{r,echo = F}
# Checks if content of a list contains entries with a ';'. If yes, then the maximum number is retrieved and put into place.
get_stderr <- function(x) {
  mean <- mean(x)
  stderr <- sd(x)/sqrt(length(x))
  return(c(mean,stderr))
}
```

## overview of observations

Amount of entries split by dsRNA method and fungal lifestyle

```{r, echo=F}
addmargins(table(edit_table$fLifestyle, edit_table$fDsRNA_method))
```

Mean resistance and standard error of resistance measurements split by dsRNA method and fungal lifestyle

```{r,echo=FALSE}
lifestyles <- c("biotroph","hemibiotroph","necrotroph")
methods <- c("HIGS","SIGS","Sum")

results <- data.frame()

for(life in lifestyles){
  for(meth in methods){
    if(meth == "Sum") {
      values <- edit_table[edit_table$fLifestyle == life, 
                           "resistance"]
    } else {
      values <- edit_table[edit_table$fLifestyle == life &
                            edit_table$fDsRNA_method == meth, 
                           "resistance"]
    }
    stats <- get_stderr(values)
    results <- rbind(results, data.frame(lifestyle = life, method = meth, mean = stats[1], SE = stats[2]))
  }
}

for (meth in methods) {
  if(meth == "Sum"){
    values <- edit_table[,"resistance"]
  } else {
    values <- edit_table[edit_table$fDsRNA_method == meth,
                         "resistance"]
  }
  stats <- get_stderr(values)
  results <- rbind(results, data.frame(lifestyle = "all", method = meth, mean = stats[1], SE = stats[2]))
}
results
```

# Reproducibility info

```{r,collapse=TRUE, code}
Sys.Date()
sessionInfo()
```
````
