## Supporting files and data for "Efficiency of RNAi based gene silencing in fungi - a review and meta-analysis": 03-generate_figures.html

Meta-analyse HIGS vs SIGS


Code 

- Show All Code
- Hide All Code

### Meta-analyse HIGS vs SIGS

###### Patrick Barth

This Markdown document generates builds a regression model and
generates figures using the edited data

### Set up

#### Session information

Preparations to set up plotting

#### Load libraries

```
packages <- c("rstudioapi","ggplot2","stringr","ggpubr","colorspace","rstatix","mgcViz","patchwork","lme4","lmerTest","effects","multcomp","metafor")
new.packages <- packages[!(packages %in% installed.packages()[,"Package"])]
if(length(new.packages) > 0) install.packages(new.packages)
library(rstudioapi)
library(ggplot2)
library(stringr)
library(ggpubr)
library(colorspace)
library(rstatix)
```

```
## 
## Attaching package: 'rstatix'
```

```
## The following object is masked from 'package:stats':
## 
##     filter
```

```
library(mgcViz)
```

```
## Loading required package: mgcv
```

```
## Loading required package: nlme
```

```
## This is mgcv 1.9-1. For overview type 'help("mgcv-package")'.
```

```
## Loading required package: qgam
```

```
## Registered S3 method overwritten by 'GGally':
##   method from   
##   +.gg   ggplot2
```

```
## Registered S3 method overwritten by 'mgcViz':
##   method from  
##   +.gg   GGally
```

```
## 
## Attaching package: 'mgcViz'
```

```
## The following objects are masked from 'package:stats':
## 
##     qqline, qqnorm, qqplot
```

```
library(patchwork)
library(lme4)
```

```
## Loading required package: Matrix
```

```
## 
## Attaching package: 'lme4'
```

```
## The following object is masked from 'package:nlme':
## 
##     lmList
```

```
library(lmerTest)
```

```
## 
## Attaching package: 'lmerTest'
```

```
## The following object is masked from 'package:lme4':
## 
##     lmer
```

```
## The following object is masked from 'package:stats':
## 
##     step
```

```
library(effects)
```

```
## Loading required package: carData
```

```
## lattice theme set by effectsTheme()
## See ?effectsTheme for details.
```

```
library(multcomp)
```

```
## Loading required package: mvtnorm
```

```
## Loading required package: survival
```

```
## Loading required package: TH.data
```

```
## Loading required package: MASS
```

```
## 
## Attaching package: 'MASS'
```

```
## The following object is masked from 'package:patchwork':
## 
##     area
```

```
## The following object is masked from 'package:rstatix':
## 
##     select
```

```
## 
## Attaching package: 'TH.data'
```

```
## The following object is masked from 'package:MASS':
## 
##     geyser
```

```
library(metafor) # Maybe remove if we continue with lme4
```

```
## Loading required package: metadat
```

```
## Loading required package: numDeriv
```

```
## 
## Loading the 'metafor' package (version 4.8-0). For an
## introduction to the package please type: help(metafor)
```

```
rm(packages,new.packages)
```

#### functions

##### Get asteriks

```
get_asteriks <- function(p){
  ast <- ifelse(p < 0.001, "***",
                ifelse(p < 0.01, "**",
                       ifelse(p < 0.05,"*",
                              ifelse(p < 0.1, ".", "n.s."))))
  return(ast)
}
```

#### Read data

```
load("../data/data_edited.RData")
```

### Meta-analysis

Our goal was to analyze the data with a random-effect meta-analysis,
which assumes that

\(y ~ x\_1 ... x\_n + \epsilon\_{measurement}
+ \epsilon\_{experiment}\)

where \(\epsilon\_{measurement}\) is
the experimental uncertainty and \(\epsilon\_{experiment}\) is an unknown
systematic effect per experiment. A common method to implement this
model is the metafor package in R. However, this package does not
support a number of downstream functions that we wanted to use,
including ANOVA. We therefore decide to implement the model in lme4,
which allows to specify nearly the same model, with one difference,
which is that \(\epsilon\_{measurement}\) is specified via
the weights argument, which means that the model uses the values up to a
constant. In other words, there is an additional df in the relative
weighting of the two errors. To check robustness to these assumptions,
we fit the model both in lme4 and in metafor. As results were very
similar, we proceeded with the lme4 model.

#### Build lme4 model

```
set.seed(123)
min_SE <- 0.001
mod.lme <- lmerTest::lmer(resistance ~ fDsRNA_method * ( fLifestyle + fFungus_class + sDsRNA_length + sNumber_of_constructs + sNumber_of_targets + sTarget_mean ) + fFormulation + (1 | id ),
                      weights = 1/((standard_error_imputed + min_SE)^2),
                      data = edit_table,
                      control = lmerControl(check.nobs.vs.nlev = "ignore",
                                            check.nobs.vs.rankZ = "ignore",
                                            check.nobs.vs.nRE = "ignore"))

mod.lme.summary <- summary(mod.lme)
mod.lme.summary
```

```
## Linear mixed model fit by REML. t-tests use Satterthwaite's method [
## lmerModLmerTest]
## Formula: 
## resistance ~ fDsRNA_method * (fLifestyle + fFungus_class + sDsRNA_length +  
##     sNumber_of_constructs + sNumber_of_targets + sTarget_mean) +  
##     fFormulation + (1 | id)
##    Data: edit_table
## Weights: 1/((standard_error_imputed + min_SE)^2)
## Control: 
## lmerControl(check.nobs.vs.nlev = "ignore", check.nobs.vs.rankZ = "ignore",  
##     check.nobs.vs.nRE = "ignore")
## 
## REML criterion at convergence: 2971.4
## 
## Scaled residuals: 
##        Min         1Q     Median         3Q        Max 
## -0.0030021 -0.0001197  0.0000046  0.0000749  0.0175941 
## 
## Random effects:
##  Groups   Name        Variance  Std.Dev.
##  id       (Intercept) 452.28085 21.26690
##  Residual               0.00232  0.04817
## Number of obs: 343, groups:  id, 343
## 
## Fixed effects:
##                                             Estimate Std. Error       df
## (Intercept)                                  59.1792     6.7559   7.3886
## fDsRNA_methodSIGS                            15.5736     5.1396  39.6196
## fLifestylehemibiotroph                       12.0542     4.4671  28.2031
## fLifestylenecrotroph                          9.6372     4.0522  22.5046
## fFungus_classBasidiomycet                     5.2595     4.3016 282.8577
## fFungus_classOomycet                         19.3317     6.6646 246.5425
## sDsRNA_length                                -0.9624     1.9483  58.3803
## sNumber_of_constructs                         1.5542     1.8172  22.7732
## sNumber_of_targets                            3.1078     2.4518  18.1003
## sTarget_mean                                  5.0257     1.6606  19.5937
## fFormulationnaked_dsRNA                      -9.1755     5.9258  27.5302
## fFormulationunknown                          -7.1055     6.1142  79.5002
## fDsRNA_methodSIGS:fLifestylehemibiotroph    -18.8192     8.7106 204.8271
## fDsRNA_methodSIGS:fLifestylenecrotroph      -14.2877     6.3097 109.9931
## fDsRNA_methodSIGS:fFungus_classBasidiomycet  -9.3782     8.6007 313.7867
## fDsRNA_methodSIGS:fFungus_classOomycet      -12.8725     8.9291 284.7194
## fDsRNA_methodSIGS:sDsRNA_length              -0.7644     2.6073 184.3819
## fDsRNA_methodSIGS:sNumber_of_constructs      -1.0951     2.7023  92.3067
## fDsRNA_methodSIGS:sNumber_of_targets         -3.0490     2.9563  60.6691
## fDsRNA_methodSIGS:sTarget_mean               -6.4014     2.4867 194.6268
##                                             t value Pr(>|t|)    
## (Intercept)                                   8.760 3.68e-05 ***
## fDsRNA_methodSIGS                             3.030  0.00429 ** 
## fLifestylehemibiotroph                        2.698  0.01163 *  
## fLifestylenecrotroph                          2.378  0.02629 *  
## fFungus_classBasidiomycet                     1.223  0.22247    
## fFungus_classOomycet                          2.901  0.00406 ** 
## sDsRNA_length                                -0.494  0.62319    
## sNumber_of_constructs                         0.855  0.40130    
## sNumber_of_targets                            1.268  0.22102    
## sTarget_mean                                  3.026  0.00678 ** 
## fFormulationnaked_dsRNA                      -1.548  0.13295    
## fFormulationunknown                          -1.162  0.24866    
## fDsRNA_methodSIGS:fLifestylehemibiotroph     -2.160  0.03190 *  
## fDsRNA_methodSIGS:fLifestylenecrotroph       -2.264  0.02551 *  
## fDsRNA_methodSIGS:fFungus_classBasidiomycet  -1.090  0.27637    
## fDsRNA_methodSIGS:fFungus_classOomycet       -1.442  0.15051    
## fDsRNA_methodSIGS:sDsRNA_length              -0.293  0.76971    
## fDsRNA_methodSIGS:sNumber_of_constructs      -0.405  0.68624    
## fDsRNA_methodSIGS:sNumber_of_targets         -1.031  0.30648    
## fDsRNA_methodSIGS:sTarget_mean               -2.574  0.01079 *  
## ---
## Signif. codes:  0 '***' 0.001 '**' 0.01 '*' 0.05 '.' 0.1 ' ' 1
```

```
## 
## Correlation matrix not shown by default, as p = 20 > 12.
## Use print(x, correlation=TRUE)  or
##     vcov(x)        if you need it
```

```
# comparison to weighted regression
# results are different, but that is what we expect, because the RE metaregression is different and preferable 

# mod.lme <- lm(resistance ~ fDsRNA_method * ( fLifestyle + fFungus_class + sDsRNA_length + sNumber_of_constructs + sNumber_of_targets + sTarget_mean ) + fFormulation ,
#                       weights = 1/((standard_error_imputed + min_SE)^2),
#                       data = edit_table)
# 
# mod.lme.summary <- summary(mod.lme)
# mod.lme.summary
```

#### Calculate ANOVA

```
anovaResults <- lmerTest:::anova.lmerModLmerTest(mod.lme, 
                                                 type = "II")
anovaResults
```

```
## Type II Analysis of Variance Table with Satterthwaite's method
##                                        Sum Sq   Mean Sq NumDF   DenDF F value
## fDsRNA_method                       0.0055122 0.0055122     1 185.360  2.3755
## fLifestyle                          0.0054712 0.0027356     2  64.731  1.1789
## fFungus_class                       0.0176200 0.0088100     2 310.486  3.7967
## sDsRNA_length                       0.0026059 0.0026059     1 149.348  1.1230
## sNumber_of_constructs               0.0014444 0.0014444     1  50.132  0.6225
## sNumber_of_targets                  0.0012671 0.0012671     1  57.753  0.5461
## sTarget_mean                        0.0071946 0.0071946     1  28.646  3.1006
## fFormulation                        0.0058936 0.0029468     2  52.643  1.2699
## fDsRNA_method:fLifestyle            0.0177777 0.0088889     2 177.315  3.8307
## fDsRNA_method:fFungus_class         0.0066175 0.0033088     2 292.233  1.4259
## fDsRNA_method:sDsRNA_length         0.0001995 0.0001995     1 184.382  0.0860
## fDsRNA_method:sNumber_of_constructs 0.0003811 0.0003811     1  92.307  0.1642
## fDsRNA_method:sNumber_of_targets    0.0024681 0.0024681     1  60.669  1.0637
## fDsRNA_method:sTarget_mean          0.0153771 0.0153771     1 194.627  6.6268
##                                      Pr(>F)  
## fDsRNA_method                       0.12496  
## fLifestyle                          0.31413  
## fFungus_class                       0.02349 *
## sDsRNA_length                       0.29098  
## sNumber_of_constructs               0.43385  
## sNumber_of_targets                  0.46292  
## sTarget_mean                        0.08894 .
## fFormulation                        0.28931  
## fDsRNA_method:fLifestyle            0.02351 *
## fDsRNA_method:fFungus_class         0.24195  
## fDsRNA_method:sDsRNA_length         0.76971  
## fDsRNA_method:sNumber_of_constructs 0.68624  
## fDsRNA_method:sNumber_of_targets    0.30648  
## fDsRNA_method:sTarget_mean          0.01079 *
## ---
## Signif. codes:  0 '***' 0.001 '**' 0.01 '*' 0.05 '.' 0.1 ' ' 1
```

#### Calculate R squared

The total R2 of the fixed effects in the model is 0.1201408

```
MuMIn::r.squaredGLMM(mod.lme)
```

```
## Registered S3 method overwritten by 'MuMIn':
##   method        from 
##   nobs.multinom broom
```

```
##            R2m       R2c
## [1,] 0.1201408 0.9999955
```

Note: the R2c is fixed + random, but this is not relevant in this
case because we use the (1|ID) RE as a random error, and thus it is
expected that fixed + random is approximately 1.

To calculate the R2 of the separate fixed effects, we first calculate
the relative SumSq (normalized to 1) of the fixed effects as displayed
by the lmerTest ANOVA table, and the multiply them with total fixed
effect R2 calculated by the r.squaredGLMM function of the MuMIn
package

```
out = data.frame(variable = rownames(anovaResults), 
                 RelativeSumSq = round(anovaResults$`Sum Sq` / sum(anovaResults$`Sum Sq`), digits = 3))
out$R2 = out$RelativeSumSq * MuMIn::r.squaredGLMM(mod.lme)[1]
out
```

```
##                               variable RelativeSumSq           R2
## 1                        fDsRNA_method         0.061 0.0073285875
## 2                           fLifestyle         0.061 0.0073285875
## 3                        fFungus_class         0.196 0.0235475927
## 4                        sDsRNA_length         0.029 0.0034840826
## 5                sNumber_of_constructs         0.016 0.0019222525
## 6                   sNumber_of_targets         0.014 0.0016819709
## 7                         sTarget_mean         0.080 0.0096112623
## 8                         fFormulation         0.066 0.0079292914
## 9             fDsRNA_method:fLifestyle         0.198 0.0237878742
## 10         fDsRNA_method:fFungus_class         0.074 0.0088904176
## 11         fDsRNA_method:sDsRNA_length         0.002 0.0002402816
## 12 fDsRNA_method:sNumber_of_constructs         0.004 0.0004805631
## 13    fDsRNA_method:sNumber_of_targets         0.027 0.0032438010
## 14          fDsRNA_method:sTarget_mean         0.171 0.0205440732
```

### Pairwise comparison plots with marginal means to show significance of differences

Pairwise comparisons are performed to analyse differences within
single predictors. We are mostly interested in whether predictors have
significant different impacts in HIGS and SIGS. Marginal means are used
to isolate the effect of single predictors using the emmeans
package.

#### Effectplot dsRNA method split by lifestyle

Pairwise comparison between HIGS and SIGS divided by the fungal
lifestyles (biotroph, hemibiotroph and necrotroph). Additionally, a plot
is generated depicting the difference between HIGS and SIGS with all
fungal lifestyles combined.

```
emm<- emmeans::emmeans(mod.lme,
                       ~ fDsRNA_method)
```

```
## NOTE: Results may be misleading due to involvement in interactions
```

```
emm_pairs <- pairs(emm, simple = "each")
emm_pairs
```

```
##  contrast    estimate   SE  df t.ratio p.value
##  HIGS - SIGS     2.88 4.36 323   0.660  0.5097
## 
## Results are averaged over the levels of: fLifestyle, fFungus_class, fFormulation 
## Degrees-of-freedom method: kenward-roger
```

```
emm_df <- as.data.frame(emm)
emm_df$fLifestyle <- "all"
emm_df$emmean
```

```
## [1] 69.17974 66.30086
```

```
emm <- emmeans::emmeans(mod.lme,
                       ~ fDsRNA_method * fLifestyle)
emm_pairs <- pairs(emm, simple = "each")
emm_pairs
```

```
## $`simple contrasts for fDsRNA_method`
## fLifestyle = biotroph:
##  contrast    estimate   SE  df t.ratio p.value
##  HIGS - SIGS    -8.16 4.89 323  -1.667  0.0964
## 
## fLifestyle = hemibiotroph:
##  contrast    estimate   SE  df t.ratio p.value
##  HIGS - SIGS    10.66 8.38 323   1.273  0.2040
## 
## fLifestyle = necrotroph:
##  contrast    estimate   SE  df t.ratio p.value
##  HIGS - SIGS     6.13 5.46 323   1.122  0.2627
## 
## Results are averaged over the levels of: fFungus_class, fFormulation 
## Degrees-of-freedom method: kenward-roger 
## 
## $`simple contrasts for fLifestyle`
## fDsRNA_method = HIGS:
##  contrast                  estimate   SE  df t.ratio p.value
##  biotroph - hemibiotroph     -12.05 4.47 323  -2.698  0.0200
##  biotroph - necrotroph        -9.64 4.05 323  -2.378  0.0471
##  hemibiotroph - necrotroph     2.42 4.13 323   0.586  0.8280
## 
## fDsRNA_method = SIGS:
##  contrast                  estimate   SE  df t.ratio p.value
##  biotroph - hemibiotroph       6.76 7.46 323   0.907  0.6365
##  biotroph - necrotroph         4.65 4.84 323   0.960  0.6025
##  hemibiotroph - necrotroph    -2.11 8.23 323  -0.257  0.9643
## 
## Results are averaged over the levels of: fFungus_class, fFormulation 
## Degrees-of-freedom method: kenward-roger 
## P value adjustment: tukey method for comparing a family of 3 estimates
```

```
emm_df <- rbind(emm_df,as.data.frame(emm))
emm_df$emmean
```

```
## [1] 69.17974 66.30086 61.94927 70.10602 74.00351 63.34110 71.58645 65.45548
```

```
pvalues <- data.frame(fLifestyle = c('all','biotroph','hemibiotroph','necrotroph'),
                      group1 = c('HIGS','HIGS','HIGS','HIGS'),
                      group2 = c('SIGS','SIGS','SIGS','SIGS'),
                      p = c(0.5097,0.0964,0.2040,0.2627))
pvalues$pAst <- apply(pvalues, 1, function(x) { get_asteriks(x['p']) })

# Double all raw values with the difference that half will be assigned to the combined lifestyle value "all"
raw_values <- attr(mod.lme, "frame")
raw_values$fLifestyle <- as.character(raw_values$fLifestyle)

raw_values_tmp <- attr(mod.lme, "frame")
raw_values_tmp$fLifestyle <- "all"
raw_values_extended <- rbind(raw_values,raw_values_tmp)

p <- ggplot(emm_df) +
  geom_violin(data = raw_values_extended, 
              aes(x = fDsRNA_method,
                  y = resistance),
              color = "gray",
              scale = "width",
              linewidth = 1.2,
              bw = 1.5) +
  geom_point(data = raw_values_extended,
             aes(x = fDsRNA_method,
                 y = resistance,
                 color = fDsRNA_method),
             position = position_jitter(width = 0.25, seed = 1),
             shape = 1,
             color = "black",
             alpha = 1,
             size = 2) +
  geom_errorbar(aes(x=fDsRNA_method,
                    ymin = lower.CL,
                    ymax = upper.CL),
                color = "black",
                width = 0.1,
                linewidth = 1)  +
    geom_point(aes(x = fDsRNA_method,
                 y = emmean),
             shape = 19,
             color = "black",
             alpha = 1,
             size = 2) +
    geom_line(aes(x=fDsRNA_method,
                 y=emmean,
                 group = fLifestyle),
              color = "black",
              linewidth = 0.5) +
  facet_wrap(~ fLifestyle,
             nrow = 1) +
  scale_y_continuous(breaks = c(-25,0,25,50,75,100),
                     limits = c(-30,115)) +
  theme(legend.position = 'none',
        panel.background = element_rect(fill = "white",
                                        color = NA),
        panel.grid.major.x = element_blank(),
        panel.grid.major.y = element_line(color = "grey",
                                          linewidth = 0.1),
        axis.ticks = element_blank(),
        axis.text = element_text(size = 12),
        axis.text.y = element_text(face = 'bold'),
        axis.title.y = element_text(size = 15),
        strip.text = element_text(size = 15)) +
  labs(x = NULL,
       y = "Resistance (%)") +
  stat_pvalue_manual(pvalues,
                    label = "{pAst}",
                    y.position = c(104,104,104,104),
                    label.size = 5,
                    tip.length = 0.001)

p
```

```
# ggsave(plot = p,
#        filename = paste(base_directory,'/../plots/','dsRNA_method_by_lifestyle.svg',sep = ''),
#        device = grDevices::svg,
#        width = 2000,
#        height = 3000,
#        units = 'px')
# # tmp since google docs does not allow the inclusion of svg
#ggsave(plot = p,
#        filename = paste('../plots/','dsRNA_method_by_lifestyle.png',sep = ''),
#        device = png,
#        width = 3000,
#        height = 2000,
#        units = 'px')
```

#### Effectplot lifestyle

Pairwise comparison of fungal lifestyles split by HIGS and SIGS

```
emm<- emmeans::emmeans(mod.lme,
                       ~ fLifestyle * fDsRNA_method)
emm_pairs <- pairs(emm, simple = "each")
emm_pairs
```

```
## $`simple contrasts for fLifestyle`
## fDsRNA_method = HIGS:
##  contrast                  estimate   SE  df t.ratio p.value
##  biotroph - hemibiotroph     -12.05 4.47 323  -2.698  0.0200
##  biotroph - necrotroph        -9.64 4.05 323  -2.378  0.0471
##  hemibiotroph - necrotroph     2.42 4.13 323   0.586  0.8280
## 
## fDsRNA_method = SIGS:
##  contrast                  estimate   SE  df t.ratio p.value
##  biotroph - hemibiotroph       6.76 7.46 323   0.907  0.6365
##  biotroph - necrotroph         4.65 4.84 323   0.960  0.6025
##  hemibiotroph - necrotroph    -2.11 8.23 323  -0.257  0.9643
## 
## Results are averaged over the levels of: fFungus_class, fFormulation 
## Degrees-of-freedom method: kenward-roger 
## P value adjustment: tukey method for comparing a family of 3 estimates 
## 
## $`simple contrasts for fDsRNA_method`
## fLifestyle = biotroph:
##  contrast    estimate   SE  df t.ratio p.value
##  HIGS - SIGS    -8.16 4.89 323  -1.667  0.0964
## 
## fLifestyle = hemibiotroph:
##  contrast    estimate   SE  df t.ratio p.value
##  HIGS - SIGS    10.66 8.38 323   1.273  0.2040
## 
## fLifestyle = necrotroph:
##  contrast    estimate   SE  df t.ratio p.value
##  HIGS - SIGS     6.13 5.46 323   1.122  0.2627
## 
## Results are averaged over the levels of: fFungus_class, fFormulation 
## Degrees-of-freedom method: kenward-roger
```

```
emm_df <- as.data.frame(emm)
emm_df$emmean
```

```
## [1] 61.94927 74.00351 71.58645 70.10602 63.34110 65.45548
```

```
pvalues <- data.frame(fDsRNA_method = c('HIGS','HIGS','HIGS','SIGS','SIGS','SIGS'),
                      group1 = c('biotroph','biotroph','hemibiotroph','biotroph','biotroph','hemibiotroph'),
                      group2 = c('hemibiotroph','necrotroph','necrotroph','hemibiotroph','necrotroph','necrotroph'),
                      p = c(0.0200,0.0471,0.8280,0.6365,0.6025,0.9643))

pvalues$pAst <- apply(pvalues, 1, function(x) { get_asteriks(x['p']) })

raw_values <- attr(mod.lme, "frame")

p <- ggplot(emm_df) +
  geom_violin(data = raw_values, 
              aes(x = fLifestyle,
                  y = resistance),
              color = "gray",
              scale = "width",
              linewidth = 1.2,
              bw = 1.5) +
  geom_point(data = raw_values,
             aes(x = fLifestyle,
                 y = resistance,
                 color = fLifestyle),
             position = position_jitter(width = 0.25, seed = 1),
             shape = 1,
             color = "black",
             alpha = 1,
             size = 2) +
  geom_errorbar(aes(x=fLifestyle,
                    ymin = lower.CL,
                    ymax = upper.CL),
                color = "black",
                width = 0.1,
                linewidth = 1) +
    geom_point(aes(x = fLifestyle,
                 y = emmean),
             shape = 19,
             color = "black",
             alpha = 1,
             size = 2) +
    geom_line(aes(x=fLifestyle,
                 y=emmean,
                 group = fDsRNA_method),
              color = "black",
              linewidth = 0.5) +
  facet_wrap(~ fDsRNA_method) +
  scale_y_continuous(breaks = c(-25,0,25,50,75,100),
                     limits = c(-30,115)) +
  theme(legend.position = 'none',
        panel.background = element_rect(fill = "white",
                                        color = NA),
        panel.grid.major.x = element_blank(),
        panel.grid.major.y = element_line(color = "grey",
                                          linewidth = 0.1),
        axis.ticks = element_blank(),
        axis.text = element_text(size = 12),
        axis.text.y = element_text(face = 'bold'),
        axis.title.y = element_text(size = 15),
        strip.text = element_text(size = 15)) +
  labs(x = NULL,
       y = "Resistance (%)") +
  stat_pvalue_manual(pvalues,
                    label = "{pAst}",
                    y.position = c(104,108,112),
                    label.size = 4,
                    tip.length = 0.001)


p
```

```
#ggsave(plot = p,
#       filename = paste(base_directory,'/../plots/','lifestyle_HIGS_vs_SIGS.svg',sep = ''),
#       device = grDevices::svg,
#       width = 3000,
#       height = 3000,
#       units = 'px')
# tmp since google docs does not allow the inclusion of svg
#ggsave(plot = p,
#      filename = paste('../plots/','lifestyle_HIGS_vs_SIGS.png',sep = ''),
#      device = png,
#      width = 3000,
#      height = 2000,
#      units = 'px')
```

#### Effectplot formulation

Pairwise comparison of samples using formulation and samples using
naked dsRNA. A third group “unknown” is added representing samples that
had no description whether a formulation was used or not. All depicted
samples are SIGS experiments. Additionally, all samples using naked RNAs
are analyzed to see if they have different effects depending on the
fungal lifestyle.

```
hjust <- 0
emm <- emmeans::emmeans(mod.lme,
                       ~ fFormulation,
                       at = list(fDsRNA_method = "SIGS"))
emm_pairs <- pairs(emm, simple = "each")
emm_pairs
```

```
##  contrast                  estimate   SE  df t.ratio p.value
##  formulation - naked_dsRNA     9.18 5.93 323   1.548  0.2698
##  formulation - unknown         7.11 6.11 323   1.162  0.4767
##  naked_dsRNA - unknown        -2.07 3.07 323  -0.674  0.7785
## 
## Results are averaged over the levels of: fLifestyle, fFungus_class 
## Degrees-of-freedom method: kenward-roger 
## P value adjustment: tukey method for comparing a family of 3 estimates
```

```
emm_df <- as.data.frame(emm)

pvalues <- data.frame(group1 = c('formulation','formulation','naked_dsRNA'),
                      group2 = c('naked_dsRNA','unknown','unknown'),
                      p = c(0.2698,0.4767,0.7785))
pvalues$pAst <- apply(pvalues, 1, function(x) { get_asteriks(x['p']) })

raw_values <- attr(mod.lme, "frame")
raw_values <- raw_values[raw_values$fDsRNA_method == "SIGS",]

p1 <- ggplot(emm_df) +
  geom_violin(data = raw_values, 
              aes(x = fFormulation,
                  y = resistance),
              color = "gray",
              scale = "width",
              linewidth = 1.2,
              bw = 1.5) +
  geom_point(data = raw_values,
             aes(x = fFormulation,
                 y = resistance,
                 color = fFormulation),
             position = position_jitter(width = 0.25, seed = 1),
             shape = 1,
             color = "black",
             alpha = 1,
             size = 2) +
  geom_errorbar(aes(x=fFormulation,
                    ymin = lower.CL,
                    ymax = upper.CL),
                color = "black",
                width = 0.1,
                linewidth = 1)  +
    geom_point(aes(x = fFormulation,
                 y = emmean),
             shape = 19,
             color = "black",
             alpha = 1,
             size = 2) +
    geom_line(aes(x=fFormulation,
                 y=emmean,
                 group = 1),
              color = "black",
              linewidth = 0.5) +
  scale_y_continuous(breaks = c(-25,0,25,50,75,100),
                     limits = c(-30,115)) +
  theme(legend.position = 'none',
        panel.background = element_rect(fill = "white",
                                        color = NA),
        panel.grid.major.x = element_blank(),
        panel.grid.major.y = element_line(color = "grey",
                                          linewidth = 0.1),
        axis.ticks = element_blank(),
        axis.text = element_text(size = 12),
        axis.text.y = element_text(face = 'bold'),
        axis.title.y = element_text(size = 15),
        strip.text = element_text(size = 15)) +
  labs(x = NULL,
       y = "Resistance (%)") +
  stat_pvalue_manual(pvalues,
                    label = "{pAst}",
                    y.position = c(104,109,114),
                    label.size = 5,
                    tip.length = 0.001) +
  annotate("text", x = -Inf, y = Inf, label = "A", hjust = hjust, vjust = 1.5, size = 15)

# Preparing the same plot for naked dsRNA only
emm<- emmeans::emmeans(mod.lme,
                       ~ fFormulation * fLifestyle,
                       at = list(fDsRNA_method = "SIGS",
                                 fFormulation = "naked_dsRNA"))
emm_pairs <- pairs(emm, simple = "each")
emm_pairs
```

```
## $`simple contrasts for fFormulation`
## fLifestyle = biotroph:
##  contrast  estimate SE df z.ratio p.value
##  (nothing)   nonEst NA NA      NA      NA
## 
## fLifestyle = hemibiotroph:
##  contrast  estimate SE df z.ratio p.value
##  (nothing)   nonEst NA NA      NA      NA
## 
## fLifestyle = necrotroph:
##  contrast  estimate SE df z.ratio p.value
##  (nothing)   nonEst NA NA      NA      NA
## 
## Results are averaged over the levels of: fFungus_class 
## Degrees-of-freedom method: kenward-roger 
## 
## $`simple contrasts for fLifestyle`
## fFormulation = naked_dsRNA:
##  contrast                  estimate   SE  df t.ratio p.value
##  biotroph - hemibiotroph       6.76 7.46 323   0.907  0.6365
##  biotroph - necrotroph         4.65 4.84 323   0.960  0.6025
##  hemibiotroph - necrotroph    -2.11 8.23 323  -0.257  0.9643
## 
## Results are averaged over the levels of: fFungus_class 
## Degrees-of-freedom method: kenward-roger 
## P value adjustment: tukey method for comparing a family of 3 estimates
```

```
emm_df <- as.data.frame(emm)

pvalues <- data.frame(group1 = c('biotroph','biotroph','hemibiotroph'),
                      group2 = c('hemibiotroph','necrotroph','necrotroph'),
                      p = c(0.6365,0.6025,0.9643))
pvalues$pAst <- apply(pvalues, 1, function(x) { get_asteriks(x['p']) })


raw_values <- attr(mod.lme, "frame")
raw_values <- raw_values[raw_values$fDsRNA_method == "SIGS",]
raw_values <- raw_values[raw_values$fFormulation == "naked_dsRNA",]

p2 <- ggplot(emm_df) +
  geom_violin(data = raw_values, 
              aes(x = fLifestyle,
                  y = resistance),
              color = "gray",
              scale = "width",
              linewidth = 1.2,
              bw = 1.5) +
  geom_point(data = raw_values,
             aes(x = fLifestyle,
                 y = resistance,
                 color = group),
             position = position_jitter(width = 0.25, seed = 1),
             shape = 1,
             color = "black",
             alpha = 1,
             size = 2) +
  geom_errorbar(aes(x=fLifestyle,
                    ymin = lower.CL,
                    ymax = upper.CL),
                color = "black",
                width = 0.1,
                linewidth = 1)  +
    geom_point(aes(x = fLifestyle,
                 y = emmean),
             shape = 19,
             color = "black",
             alpha = 1,
             size = 2) +
    geom_line(aes(x=fLifestyle,
                 y=emmean,
                 group = 1),
              color = "black",
              linewidth = 0.5) +
  scale_y_continuous(breaks = c(-25,0,25,50,75,100),
                     limits = c(-30,115)) +
  theme(legend.position = 'none',
        panel.background = element_rect(fill = "white",
                                        color = NA),
        panel.grid.major.x = element_blank(),
        panel.grid.major.y = element_line(color = "grey",
                                          linewidth = 0.1),
        axis.ticks = element_blank(),
        axis.text = element_text(size = 12),
        axis.text.y = element_text(face = 'bold'),
        axis.title.y = element_text(size = 15),
        strip.text = element_text(size = 15)) +
  labs(x = NULL,
       y = "Resistance (naked dsRNA)") +
  stat_pvalue_manual(pvalues,
                    label = "{pAst}",
                    y.position = c(104,109,114),
                    label.size = 5,
                    tip.length = 0.001) +
  annotate("text", x = -Inf, y = Inf, label = "B", hjust = hjust, vjust = 1.5, size = 15)

p <- p1 | p2


p
```

```
# ggsave(plot = p,
#        filename = paste(base_directory,'/../plots/','formulation_and_naked_dsRNA_by_lifestyle.svg',sep = ''),
#        device = grDevices::svg,
#        width = 3000,
#        height = 2000,
#        units = 'px')
# # tmp since google docs does not allow the inclusion of svg
#ggsave(plot = p,
#        filename = paste('../plots/','formulation_and_naked_dsRNA_by_lifestyle.png',sep = ''),
#        device = png,
#        width = 3000,
#        height = 2000,
#        units = 'px')
```

#### Effectplot dsRNA length

Analysis to see if the dsRNA length has an impact in either HIGS or
SIGS experiments

```
emm<- emmeans::emmeans(mod.lme,
                       ~ sDsRNA_length * fDsRNA_method)

emm_trends <- emmeans::emtrends(mod.lme, 
                                ~ fDsRNA_method,
                                var = "sDsRNA_length")

# Get p-values for each interaction separated
p.value_sep <- summary(emm_trends, infer = c(TRUE,TRUE))

pvalues <- data.frame(fDsRNA_method = p.value_sep$fDsRNA_method,
                      p = p.value_sep$p.value)
pvalues$pAst <- apply(pvalues, 1, function(x) { get_asteriks(x['p']) })

# Get p-value for difference between interaction
p.value_dif <- emmeans::contrast(emm_trends,
                  method = "pairwise")

pvalues_2 <- data.frame( group1 = "HIGS",
                         group2 = "SIGS",
                      xmin = 200,
                      xmax = 900,
                      p = as.data.frame(p.value_dif)$p.value)
pvalues_2$pAst <- apply(pvalues_2, 1, function(x) { get_asteriks(x['p']) })

raw_values <- attr(mod.lme, "frame")

# Generate marginal lines
all_lengths <- attr(mod.lme, "frame")$sDsRNA_length
dsRNA_length_X <- seq(min(all_lengths),
                      max(all_lengths),
                      length.out = 100)

emm <- emmeans::emmeans(mod.lme,
                        ~ sDsRNA_length * fDsRNA_method,
                        at = list(sDsRNA_length = dsRNA_length_X))
emm_df <- as.data.frame(emm)

# Prepare letters to be depicted in plots
facet_labels <- data.frame(
  fDsRNA_method = c("HIGS", "SIGS"),
  label = c("A", "B"),
  sDsRNA_length = c(min(all_lengths), min(all_lengths)),
  resistance = c(110, 110)
)


p1 <- ggplot(emm_df) +
  geom_point(data = raw_values,
             aes(x = sDsRNA_length,
                 y = resistance),
             color = "black",
             position = position_jitter(width = 0.25, seed = 1),
             shape = 1) +
    geom_line(aes(x=sDsRNA_length,
                 y=emmean,
                 group = fDsRNA_method),
            color = "black",
            linewidth = 0.5) +
  geom_ribbon(aes(x = sDsRNA_length,
                  ymin = lower.CL,
                  ymax = upper.CL),
              alpha = 0.2) +
  facet_wrap(~ fDsRNA_method) +
  scale_y_continuous(breaks = c(-25,0,25,50,75,100),
                     limits = c(-30,115)) +
  theme(legend.position = 'none',
        panel.background = element_rect(fill = "white",
                                        color = NA),
        panel.grid.major.x = element_blank(),
        panel.grid.major.y = element_line(color = "grey",
                                          linewidth = 0.1),
        axis.ticks = element_blank(),
        axis.text = element_text(size = 12),
        axis.text.y = element_text(face = 'bold'),
        axis.title.y = element_text(size = 15),
        strip.text = element_text(size = 15),
        plot.title = element_text(hjust = 0.5, size = 15, color = "black")) +
  labs(title = "dsRNA length",
       x = NULL,
       y = "Resistance (%)") +
  geom_text(data = pvalues,
            aes(x = 1,
                y = 105,
                label = paste("p = ", pAst, sep = "")),
            inherit.aes = FALSE,
            size = 5) +
  geom_text(data = facet_labels,
    aes(x = sDsRNA_length,
        y = resistance,
        label = label),
        inherit.aes = FALSE,
        size = 10)


p1
```

#### Effectplot target position

Analysis to see if the dsRNA position upon the target gene has an
impact in either HIGS or SIGS experiments

```
emm<- emmeans::emmeans(mod.lme,
                       ~ sTarget_mean * fDsRNA_method)

emm_trends <- emmeans::emtrends(mod.lme, 
                                ~ fDsRNA_method,
                                var = "sTarget_mean")

# Get p-values for each interaction separated
p.value_sep <- summary(emm_trends, infer = c(TRUE,TRUE))

pvalues <- data.frame(fDsRNA_method = p.value_sep$fDsRNA_method,
                      p = p.value_sep$p.value)
pvalues$pAst <- apply(pvalues, 1, function(x) { get_asteriks(x['p']) })

# Get p-value for difference between interaction
p.value_dif <- emmeans::contrast(emm_trends,
                  method = "pairwise")

pvalues_2 <- data.frame( group1 = "HIGS",
                         group2 = "SIGS",
                      xmin = 200,
                      xmax = 900,
                      p = as.data.frame(p.value_dif)$p.value)
pvalues_2$pAst <- apply(pvalues_2, 1, function(x) { get_asteriks(x['p']) })

raw_values <- attr(mod.lme, "frame")

# Generate marginal lines
all_lengths <- attr(mod.lme, "frame")$sTarget_mean
dsRNA_length_X <- seq(min(all_lengths),
                      max(all_lengths),
                      length.out = 100)

emm <- emmeans::emmeans(mod.lme,
                        ~ sTarget_mean * fDsRNA_method,
                        at = list(sTarget_mean = dsRNA_length_X))
emm_df <- as.data.frame(emm)

# Prepare letters to be depicted in plots
facet_labels <- data.frame(
  fDsRNA_method = c("HIGS", "SIGS"),
  label = c("C", "D"),
  sTarget_mean = c(min(all_lengths), min(all_lengths)),
  resistance = c(110, 110)
)

p2 <- ggplot(emm_df) +
  geom_point(data = raw_values,
             aes(x = sTarget_mean,
                 y = resistance),
             color = "black",
             position = position_jitter(width = 0.25, seed = 1),
             shape = 1) +
    geom_line(aes(x=sTarget_mean,
                 y=emmean,
                 group = fDsRNA_method),
            color = "black",
            linewidth = 0.5) +
  geom_ribbon(aes(x = sTarget_mean,
                  ymin = lower.CL,
                  ymax = upper.CL),
              alpha = 0.2) +
  facet_wrap(~ fDsRNA_method) +
  scale_y_continuous(breaks = c(-25,0,25,50,75,100),
                     limits = c(-30,115)) +
  theme(legend.position = 'none',
        panel.background = element_rect(fill = "white",
                                        color = NA),
        panel.grid.major.x = element_blank(),
        panel.grid.major.y = element_line(color = "grey",
                                          linewidth = 0.1),
        axis.ticks = element_blank(),
        axis.text = element_text(size = 12),
        axis.text.y = element_text(face = 'bold'),
        axis.title.y = element_text(size = 15),
        strip.text = element_text(size = 15),
        plot.title = element_text(hjust = 0.5, size = 15, color = "black")) +
  labs(title = "Target position on mRNA",
       x = NULL,
       y = "Resistance (%)") +
  geom_text(data = pvalues,
            aes(x = 0,
                y = 105,
                label = paste("p = ", pAst, sep = "")),
            inherit.aes = FALSE,
            size = 5) +
  geom_text(data = facet_labels,
    aes(x = sTarget_mean,
        y = resistance,
        label = label),
        inherit.aes = FALSE,
        size = 10)

p2
```

```
p <- p1 / p2
# ggsave(plot = p,
#        filename = paste(base_directory,'/../plots/','dsRNA_length_and_target_position.svg',sep = ''),
#        device = grDevices::svg,
#        width = 3000,
#        height = 3000,
#        units = 'px')
# # tmp since google docs does not allow the inclusion of svg
# ggsave(plot = p,
#        filename = paste('../plots/','dsRNA_length_and_target_position.png',sep = ''),
#        device = png,
#        width = 3000,
#        height = 3000,
#        units = 'px')
```

#### Effect of construct and target gene number for supplements

Analysis to see if the number of used constructs has an impact in
either HIGS or SIGS experiments

```
# Number of constructs
emm<- emmeans::emmeans(mod.lme,
                       ~ sNumber_of_constructs * fDsRNA_method)

emm_trends <- emmeans::emtrends(mod.lme, 
                                ~ fDsRNA_method,
                                var = "sNumber_of_constructs")

# Get p-values for each interaction separated
p.value_sep <- summary(emm_trends, infer = c(TRUE,TRUE))

pvalues <- data.frame(fDsRNA_method = p.value_sep$fDsRNA_method,
                      p = p.value_sep$p.value)
pvalues$pAst <- apply(pvalues, 1, function(x) { get_asteriks(x['p']) })

# Get p-value for difference between interaction
p.value_dif <- emmeans::contrast(emm_trends,
                  method = "pairwise")

pvalues_2 <- data.frame( group1 = "HIGS",
                         group2 = "SIGS",
                      xmin = 200,
                      xmax = 900,
                      p = as.data.frame(p.value_dif)$p.value)
pvalues_2$pAst <- apply(pvalues_2, 1, function(x) { get_asteriks(x['p']) })

raw_values <- attr(mod.lme, "frame")

# Generate marginal lines
all_lengths <- attr(mod.lme, "frame")$sNumber_of_constructs
number_of_constructs_X <- seq(min(all_lengths),
                              max(all_lengths),
                              length.out = 100)

emm <- emmeans::emmeans(mod.lme,
                        ~ sNumber_of_constructs * fDsRNA_method,
                        at = list(sNumber_of_constructs = number_of_constructs_X))
emm_df <- as.data.frame(emm)

# Prepare letters to be depicted in plots
facet_labels <- data.frame(
  fDsRNA_method = c("HIGS", "SIGS"),
  label = c("A", "B"),
  sNumber_of_constructs = c(min(all_lengths), min(all_lengths)),
  resistance = c(110, 110)
)

p1 <- ggplot(emm_df) +
  geom_point(data = raw_values,
             aes(x = sNumber_of_constructs,
                 y = resistance),
             color = "black",
             position = position_jitter(width = 0.25, seed = 1),
             shape = 1) +
    geom_line(aes(x=sNumber_of_constructs,
                 y=emmean,
                 group = fDsRNA_method),
            color = "black",
            linewidth = 0.5) +
  geom_ribbon(aes(x = sNumber_of_constructs,
                  ymin = lower.CL,
                  ymax = upper.CL),
              alpha = 0.2) +
  facet_wrap(~ fDsRNA_method) +
  scale_y_continuous(breaks = c(-25,0,25,50,75,100),
                     limits = c(-30,115)) +
  theme(legend.position = 'none',
        panel.background = element_rect(fill = "white",
                                        color = NA),
        panel.grid.major.x = element_blank(),
        panel.grid.major.y = element_line(color = "grey",
                                          linewidth = 0.1),
        axis.ticks = element_blank(),
        axis.text = element_text(size = 12),
        axis.text.y = element_text(face = 'bold'),
        axis.title.y = element_text(size = 15),
        strip.text = element_text(size = 15),
        plot.title = element_text(hjust = 0.5, size = 15, color = "black")) +
  labs(title = "Number of constructs",
       x = NULL,
       y = "Resistance (%)") +
  geom_text(data = pvalues,
            aes(x = 3,
                y = 105,
                label = paste("p = ", pAst, sep = "")),
            inherit.aes = FALSE,
            size = 5) +
  geom_text(data = facet_labels,
    aes(x = sNumber_of_constructs,
        y = resistance,
        label = label),
        inherit.aes = FALSE,
        size = 10)

p1
```

Analysis to see if the number of targets has an impact in either HIGS
or SIGS experiments

```
emm<- emmeans::emmeans(mod.lme,
                       ~ sNumber_of_targets * fDsRNA_method)

emm_trends <- emmeans::emtrends(mod.lme, 
                                ~ fDsRNA_method,
                                var = "sNumber_of_targets")

# Get p-values for each interaction separated
p.value_sep <- summary(emm_trends, infer = c(TRUE,TRUE))

pvalues <- data.frame(fDsRNA_method = p.value_sep$fDsRNA_method,
                      p = p.value_sep$p.value)
pvalues$pAst <- apply(pvalues, 1, function(x) { get_asteriks(x['p']) })

# Get p-value for difference between interaction
p.value_dif <- emmeans::contrast(emm_trends,
                  method = "pairwise")

pvalues_2 <- data.frame( group1 = "HIGS",
                         group2 = "SIGS",
                      xmin = 200,
                      xmax = 900,
                      p = as.data.frame(p.value_dif)$p.value)
pvalues_2$pAst <- apply(pvalues_2, 1, function(x) { get_asteriks(x['p']) })

raw_values <- attr(mod.lme, "frame")

# Generate marginal lines
all_lengths <- attr(mod.lme, "frame")$sNumber_of_targets
number_of_targets_X <- seq(min(all_lengths),
                              max(all_lengths),
                              length.out = 100)

emm <- emmeans::emmeans(mod.lme,
                        ~ sNumber_of_targets * fDsRNA_method,
                        at = list(sNumber_of_targets = number_of_targets_X))
emm_df <- as.data.frame(emm)

# Prepare letters to be depicted in plots
facet_labels <- data.frame(
  fDsRNA_method = c("HIGS", "SIGS"),
  label = c("C", "D"),
  sNumber_of_targets = c(min(all_lengths), min(all_lengths)),
  resistance = c(110, 110)
)

p2 <- ggplot(emm_df) +
  geom_point(data = raw_values,
             aes(x = sNumber_of_targets,
                 y = resistance),
             color = "black",
             position = position_jitter(width = 0.25, seed = 1),
             shape = 1) +
    geom_line(aes(x=sNumber_of_targets,
                 y=emmean,
                 group = fDsRNA_method),
            color = "black",
            linewidth = 0.5) +
  geom_ribbon(aes(x = sNumber_of_targets,
                  ymin = lower.CL,
                  ymax = upper.CL),
              alpha = 0.2) +
  facet_wrap(~ fDsRNA_method) +
  scale_y_continuous(breaks = c(-25,0,25,50,75,100),
                     limits = c(-30,115)) +
  theme(legend.position = 'none',
        panel.background = element_rect(fill = "white",
                                        color = NA),
        panel.grid.major.x = element_blank(),
        panel.grid.major.y = element_line(color = "grey",
                                          linewidth = 0.1),
        axis.ticks = element_blank(),
        axis.text = element_text(size = 12),
        axis.text.y = element_text(face = 'bold'),
        axis.title.y = element_text(size = 15),
        strip.text = element_text(size = 15),
        plot.title = element_text(hjust = 0.5, size = 15, color = "black")) +
  labs(title = "Number of targets",
       x = NULL,
       y = "Resistance (%)") +
  geom_text(data = pvalues,
            aes(x = 3,
                y = 105,
                label = paste("p = ", pAst, sep = "")),
            inherit.aes = FALSE,
            size = 5) +
  geom_text(data = facet_labels,
    aes(x = sNumber_of_targets,
        y = resistance,
        label = label),
        inherit.aes = FALSE,
        size = 10)

p2
```

```
p <- p1 / p2
# ggsave(plot = p,
#        filename = paste('../plots/','number_of_constructs_and_targets.svg',sep = ''),
#        device = grDevices::svg,
#        width = 3000,
#        height = 3000,
#        units = 'px')
# # tmp since google docs does not allow the inclusion of svg
#ggsave(plot = p,
#        filename = paste('../plots/','number_of_constructs_and_targets.png',sep = ''),
#        device = png,
#        width = 3000,
#        height = 3000,
#        units = 'px')
```

```
Sys.Date()
## [1] "2025-12-08"
sessionInfo()
## R version 4.3.3 (2024-02-29)
## Platform: x86_64-pc-linux-gnu (64-bit)
## Running under: Ubuntu 24.04.3 LTS
## 
## Matrix products: default
## BLAS:   /usr/lib/x86_64-linux-gnu/blas/libblas.so.3.12.0 
## LAPACK: /usr/lib/x86_64-linux-gnu/lapack/liblapack.so.3.12.0
## 
## locale:
##  [1] LC_CTYPE=en_US.UTF-8       LC_NUMERIC=C              
##  [3] LC_TIME=en_US.UTF-8        LC_COLLATE=en_US.UTF-8    
##  [5] LC_MONETARY=en_US.UTF-8    LC_MESSAGES=en_US.UTF-8   
##  [7] LC_PAPER=en_US.UTF-8       LC_NAME=C                 
##  [9] LC_ADDRESS=C               LC_TELEPHONE=C            
## [11] LC_MEASUREMENT=en_US.UTF-8 LC_IDENTIFICATION=C       
## 
## time zone: Europe/Berlin
## tzcode source: system (glibc)
## 
## attached base packages:
## [1] stats     graphics  grDevices utils     datasets  methods   base     
## 
## other attached packages:
##  [1] metafor_4.8-0       numDeriv_2016.8-1.1 metadat_1.4-0      
##  [4] multcomp_1.4-28     TH.data_1.1-3       MASS_7.3-60.0.1    
##  [7] survival_3.5-8      mvtnorm_1.3-3       effects_4.2-2      
## [10] carData_3.0-5       lmerTest_3.1-3      lme4_1.1-37        
## [13] Matrix_1.6-5        patchwork_1.3.0     mgcViz_0.2.0       
## [16] qgam_2.0.0          mgcv_1.9-1          nlme_3.1-168       
## [19] rstatix_0.7.2       colorspace_2.1-1    ggpubr_0.6.0       
## [22] stringr_1.5.1       ggplot2_3.5.2       rstudioapi_0.17.1  
## 
## loaded via a namespace (and not attached):
##  [1] Rdpack_2.6.4       DBI_1.2.3          gridExtra_2.3      sandwich_3.1-1    
##  [5] rlang_1.1.6        magrittr_2.0.3     matrixStats_1.5.0  compiler_4.3.3    
##  [9] vctrs_0.6.5        pkgconfig_2.0.3    fastmap_1.2.0      backports_1.5.0   
## [13] labeling_0.4.3     promises_1.3.2     rmarkdown_2.29     nloptr_2.2.1      
## [17] purrr_1.0.4        xfun_0.52          cachem_1.1.0       jsonlite_2.0.0    
## [21] later_1.4.2        broom_1.0.8        parallel_4.3.3     R6_2.6.1          
## [25] bslib_0.9.0        stringi_1.8.7      RColorBrewer_1.1-3 GGally_2.2.1      
## [29] car_3.1-3          boot_1.3-30        estimability_1.5.1 jquerylib_0.1.4   
## [33] Rcpp_1.0.14        iterators_1.0.14   knitr_1.50         zoo_1.8-14        
## [37] httpuv_1.6.16      splines_4.3.3      nnet_7.3-19        tidyselect_1.2.1  
## [41] abind_1.4-8        yaml_2.3.10        viridis_0.6.5      MuMIn_1.48.4      
## [45] doParallel_1.0.17  codetools_0.2-19   miniUI_0.1.2       lattice_0.22-5    
## [49] tibble_3.2.1       plyr_1.8.9         shiny_1.10.0       withr_3.0.2       
## [53] evaluate_1.0.3     ggstats_0.9.0      survey_4.4-2       pillar_1.10.2     
## [57] KernSmooth_2.23-22 foreach_1.5.2      stats4_4.3.3       reformulas_0.4.1  
## [61] insight_1.3.0      generics_0.1.4     mathjaxr_1.8-0     scales_1.4.0      
## [65] minqa_1.2.8        xtable_1.8-4       gamm4_0.2-6        glue_1.8.0        
## [69] emmeans_1.11.1     tools_4.3.3        ggsignif_0.6.4     grid_4.3.3        
## [73] tidyr_1.3.1        mitools_2.4        rbibutils_2.3      Formula_1.2-5     
## [77] cli_3.6.5          viridisLite_0.4.2  dplyr_1.1.4        gtable_0.3.6      
## [81] sass_0.4.10        digest_0.6.37      pbkrtest_0.5.4     farver_2.1.2      
## [85] htmltools_0.5.8.1  lifecycle_1.0.4    mime_0.13
```

#### tmp
